# RNA polymerase II condensate formation and association with Cajal and histone locus bodies in living human cells

**DOI:** 10.1101/2020.12.25.424380

**Authors:** Takashi Imada, Takeshi Shimi, Ai Kaiho, Yasushi Saeki, Hiroshi Kimura

## Abstract

In eukaryotic nuclei, a number of phase-separated nuclear bodies (NBs) are present. RNA polymerase II (Pol II) is the main player in transcription and forms large condensates in addition to localizing at numerous transcription foci. Cajal bodies (CBs) and histone locus bodies (HLBs) are NBs that are involved in transcriptional and post-transcriptional regulation of small nuclear RNA and histone genes. By live-cell imaging using human HCT116 cells, we here show that Pol II condensates (PCs) nucleated near CBs and HLBs, and the number of PCs increased during S phase concomitantly with the activation period of histone genes. Ternary PC–CB– HLB associates were formed via three pathways: nucleation of PCs and HLBs near CBs, interaction between preformed PC–HLBs with CBs, and nucleation of PCs near preformed CB– HLBs. Coilin knockout increased the co-localization rate between PCs and HLBs, whereas the number, nucleation timing, and phosphorylation status of PCs remained unchanged. Depletion of PCs did not affect CBs and HLBs. Treatment with 1,6-hexanediol revealed that PCs were more liquid-like than CBs and HLBs. Thus, PCs are dynamic structures often nucleated following the activation of gene clusters associated with other NBs. (187 words)

## 1. INTRODUCTION

Eukaryotic nuclei are highly organized structures comprising chromatin, the nuclear lamina, nucleoli, and nuclear bodies (NBs). NBs are phase-separated liquid droplets that contain specific proteins and RNAs [Uversky, 2017; Strom et al, 2019]. In humans, RNA polymerase II (Pol II) is the main player in transcription and is composed of 12 subunits. In addition to numerous small transcription foci, Pol II can form relatively large phase-separated condensates via the intrinsically disordered C-terminal domain (CTD) of the Rpb1 subunit, which is the largest subunit that acts as the catalytic subunit of Pol II [Boehning et al, 2018; Lu et al, 2018]. Human Pol II CTD consists of 52 repeats of hepta-amino acids, Tyr–Ser–Pro–Thr–Ser–Pro– Ser. The Tyr, Ser, and Thr residues can be phosphorylated, depending on the transcription status. In general, Ser5 and Ser2 become phosphorylated during the initiation and elongation of transcription, respectively [Harlen and Churchman, 2017; Maita and Nakagawa, 2020]. Thus, transcribing Pol II molecules on most of protein coding genes harbor phosphorylated Ser2 (S2P) at the CTD, but those on histone genes and small nuclear RNA (snRNA) genes are only phosphorylated at Ser5 (S5P) [Medlin et al, 2005]. Pol II on snRNA genes also harbors phosphorylated Ser7 (S7P) [Egloff et al, 2007; Egloff et al, 2012]. Phosphorylation of CTD was recently reported to regulate phase separation of Pol II clusters or condensates. Pol II with hypophosphorylated CTD forms clusters with mediators, while Pol II with hyperphosphorylated CTD is clustered with splicing factors [Guo et al, 2019]. Pol II large condensates [hereafter termed Pol II condensates (PCs) to avoid confusion with numerous smaller clusters] contain both unphosphorylated and S5P CTD forms [Schul et al, 1998; Xie and Pombo, 2006; Alm-Kristiansen et al, 2008; Nizami et al, 2010; Guglielmi et al, 2013].

PCs were reported to associate with other NBs, such as Cajal/coiled bodies (CBs) and histone locus bodies (HLBs) [Lamond and Carmo-Fonseca, 1993; Schul et al, 1998; Xie and Pombo, 2006; Alm-Kristiansen et al, 2008; Nizami et al, 2010; Guglielmi et al, 2013]. CBs are involved in transcriptional and post-transcriptional regulation of histone genes, snRNA genes, and small nucleolar RNA (snoRNA) genes via genome organization [Wang et al, 2016], small nuclear ribonucleoprotein (snRNP) biogenesis [Staněk, 2017], and telomerase RNA modification [Venteicher et al, 2009; Henriksson and Farnebo, 2015]. CBs tend to localize near sn/snoRNA genes (U1 and U2 snRNA gene arrays on chromosomes 1 and 17, respectively) and *HIST2* locus (six histone genes on chromosome 1) in human cells [Frey and Matera, 2001; Shopland et al, 2001; Nizami et al, 2010a,b; Wang et al, 2016; Duronio and Marzluff, 2017]. CBs contain various proteins, such as coilin, survival motor neuron protein (SMN) [Raimer et al, 2016], and telomerase Cajal body protein 1 (TCAB1; also known as WDR79 or WRAP53β) [Venteicher et al, 2009; Henriksson and Farnebo, 2015]. Coilin is a marker protein of CBs, and its depletion leads to loss of CBs [Machyna et al, 2015]. SMN and TCAB1 contribute to snRNP recycling and telomere maintenance, respectively [Venteicher et al, 2009; Henriksson and Farnebo, 2015; Raimer et al, 2016]. Diverse biological functions of coilin or CBs have been observed during development and reproduction. Coilin-knockout (KO) mice show semi-embryonic lethality and are defective in fertility and fecundity [Tucker et al, 2001; Walker et al, 2009], and coilin-knockdown (KD) zebrafish were embryonic lethal due to insufficient snRNP production [Strzelecka et al, 2010]. At the cellular level, coilin-KO mouse and coilin-KD human cells are viable but show impaired splicing efficiency and fidelity, decreased transcription of histone and sn/snoRNA genes, and decreased cell proliferation [Tucker et al, 2001; Lemm et al, 2006; Whittom et al, 2008; Wang et al, 2016]. Coilin is considered to play a role as glue to co-assemble many proteins and RNAs into CBs and to facilitate RNP biogenesis [Klingauf et al, 2006].

The HLB is another type of NB that localizes near replication-dependent histone gene loci, such as *HIST1* (55 histone genes on chromosome 6) and *HIST2* (6 histone genes on chromosome 1) in humans [Marzluff, 2002; Nizami et al, 2010a,b; Wang et al, 2016; Duronio and Marzluff, 2017]. HLBs are often found adjacent to CBs, consistent with the association between CBs and *HIST2* [Nizami et al, 2010a,b; Duronio and Marzluff, 2017]. HLBs contain proteins essential for both the initiation of histone gene transcription and 3′ cleavage of non-polyadenylated histone mRNAs, including an HLB marker protein, nuclear protein of the ataxia telangiectasia mutated locus (NPAT) [Fruscio et al, 1997; Ma et al, 2000; Ye et al, 2003; Ghule et al, 2008; Romeo and Schümperli, 2016]. Disruption of the *NPAT* gene results in early embryonic lethality in mice and halts cell proliferation due to G_0_/G_1_ arrest [Di Fruscio et al, 1997; Ma et al, 2000; Ye et al, 2003].

Although the liquid droplet nature of NBs has recently been studied intensively [Carmo-Fonseca and Rino, 2011; Shevtsov and Dundr, 2011; Staněk and Fox 2017; Sawyer et al, 2019], it remains unclear how different NBs associate with each other in living cells. The present study, time-lapse imaging was employed to examine how three NBs (PCs, CBs, and HLBs) are formed and interact with each other in living human cells. PCs were often nucleated at the beginning of S phase near CBs or HLBs, with a preferential association with HLBs. PC– CB–HLB ternary associates were formed via three different pathways. Although CBs were not essential for PC formation, the association of PCs with other NBs was affected. PCs were more sensitive to 1,6-hexanediol (1,6-HD) than CBs and HLBs. Our data combined with previous findings suggest that PCs are dynamic structures, and their nucleation is initiated during activation of gene clusters, facilitated by other NBs.

## 2. RESULTS

### 2.1. Frequent emergence of PCs at entry of S phase in living cells

To examine when and where PCs are formed in living cells, we knocked-in the enhanced green fluorescent protein (EGFP) into both alleles of Rpb1 gene in human colon cancer HCT116 cells so that EGFP-Rpb1 is expressed under the own promotor (Fig. 1A). Western blot analysis confirmed that EGFP–Rpb1 was expressed in knock-in HCT116 cells (referred as KI cells) at a similar level to that of endogenous Rpb1 in wild type cells (Fig. 1B). In living KI cells, EGFP–Rpb1 autofluorescence was concentrated in several (average 5.4 ± 2.2; range 0–10; Table S1) large foci (Pol II condensates, or PCs) with a maximum Feret diameter of ∼0.7 μm (Fig. 1C), as previously reported [Steurer et al, 2018]. Immunofluorescence using phosphorylation-specific antibodies showed that PCs contained S5P and S7P, but not S2P and T4P, on Pol II CTD (Fig. 1D), which was consistent with observations using parental HCT116 cells (Fig. S1) as well as previous reports [Xie and Pombo, 2006].

**Figure 1.**
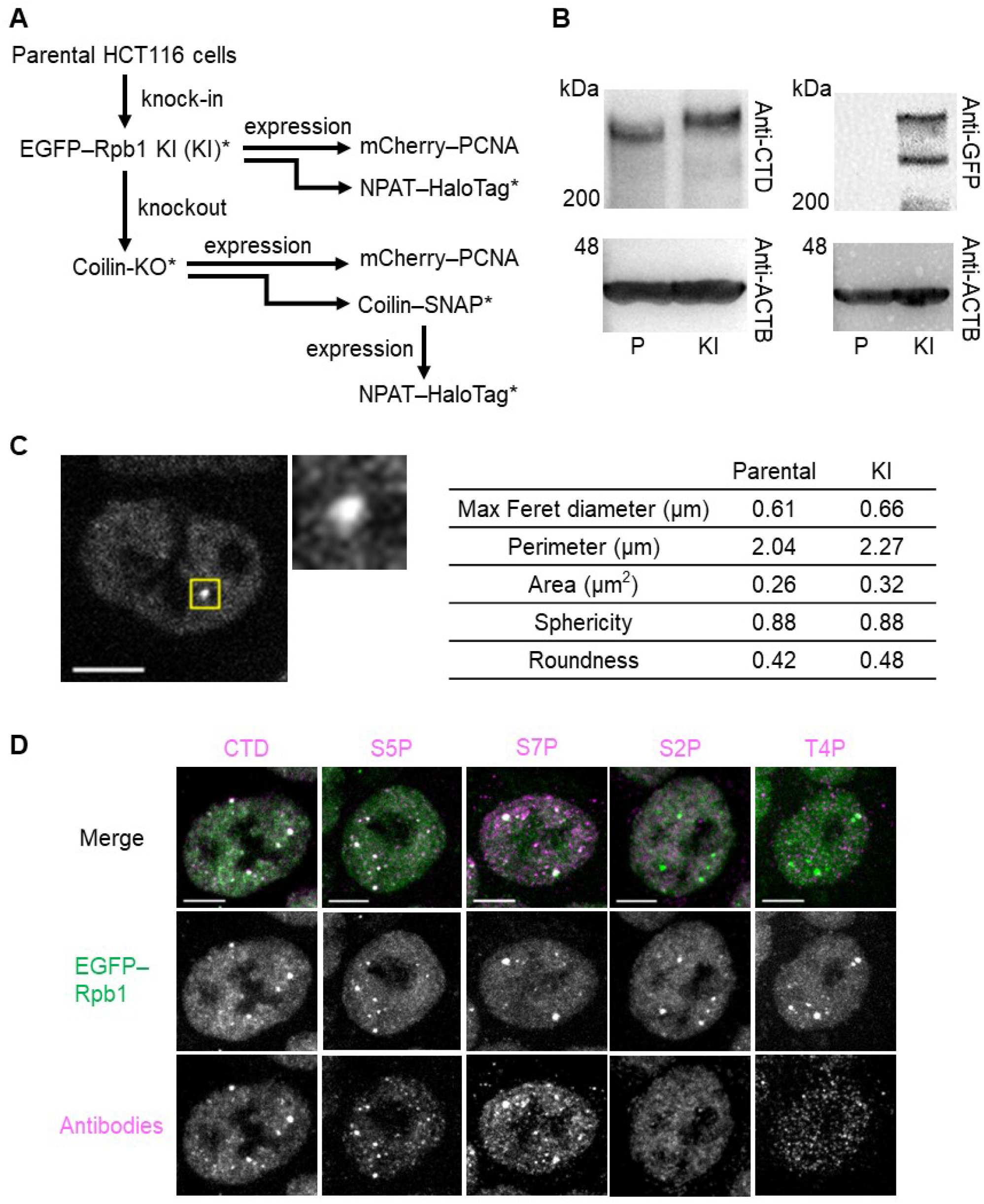
Generation of EGFP–Rpb1 KI cells and characterization of PCs. **(A)** Summary of the cell lines used in the present study. EGFP–Rpb1 KI (KI) is a cell line in which EGFP is knocked-in into both alleles and all Rpb1 is EGFP-tagged. Asterisks indicate cell lines obtained via single-cell cloning. **(B)** Western blots. Whole cell lysates prepared from parental HCT116 (P) and KI (KI) cells were separated and probed with antibodies against Pol II CTD, GFP, and beta-actin (ACTB). **(C)** EGFP–Rpb1 in living KI cells. Single confocal section of EGFP–Rpb1 and magnified view of a Pol II condensate (PC) are shown. The table (right) shows the properties of PCs in parental HCT116 and KI cells, detected by immunofluorescence using anti-CTD and EGFP–Rpb1, respectively. Parameters (means of 50 cells) were determined by intensity thresholding and size constraint (see Experimental Procedures section). The number of CBs and HLBs in KI cells (2.8 ± 1.3 and 3.3 ± 1.1, respectively; n = 100 cells; Fig. S2A; Table S1) were similar to those in parental HCT116 cells (2.8 ± 1.2 and 3.5 ± 1.3, respectively; n = 100 cells; Fig. S2B; Table S1). **(D)** Immunofluorescence images of EGFP–Rpb1 (green) in KI cells with antibodies against Pol II CTD, S2P, T4P, S5P, or S7P, and Cy5-conjugated anti-mouse secondary antibodies (magenta). Max-Intensity-Projection (MIP) images of Z-stack confocal sections at 0.30 μm intervals (31– 34 stacks) to cover the entire nucleus are shown. Scale bars, 5 μm.

We expressed mCherry–PCNA as a marker of replication foci during S phase [Leonhardt et al, 2000] in KI cells to examine the cell cycle dynamics of PCs. Fig. 2A shows a typical example representing the dynamics of EGFP–Rpb1 and mCherry–PCNA during the cell cycle (Movie S1). A few bright PCs were observed from G_2_ to M phase (elapse time, h:min; 0, 2:40, and 3:00). After the mitosis, PCs were rarely observed during G_1_ phase (4:10 and 7:10). The number of PCs increased when the cells entered S phase (7:40, at which mCherry–PCNA was concentrated in many replication foci). Most PCs persisted until the mid to late S phase (13:50 and 16:50) and the number decreased during the G_2_ to the next G_1_ phase (9:50, 21:50, 22:30, and 26:20). Analysis of 100 cells revealed that the number of PCs per nucleus increased from 0.3 ± 0.5 in G_1_ phase (0 in 69% of cells) to 3.0 ± 1.0 in early S phase (3 in 46% of cells) (Fig. 2B). It was plausible that PCs appeared in G_1_ and S phase are associated with CBs and HLBs that are involved in gene expression of snRNA (expressed in G_1_ phase and enhanced in S phase) and histone genes (expressed in S phase) [Faresse et al, 2012; Wang et al, 2016; Marzluff and Koreski, 2017]. This prompted us to investigate the relationship between PCs, CBs, and HLBs in living cells.

**Figure 2.**
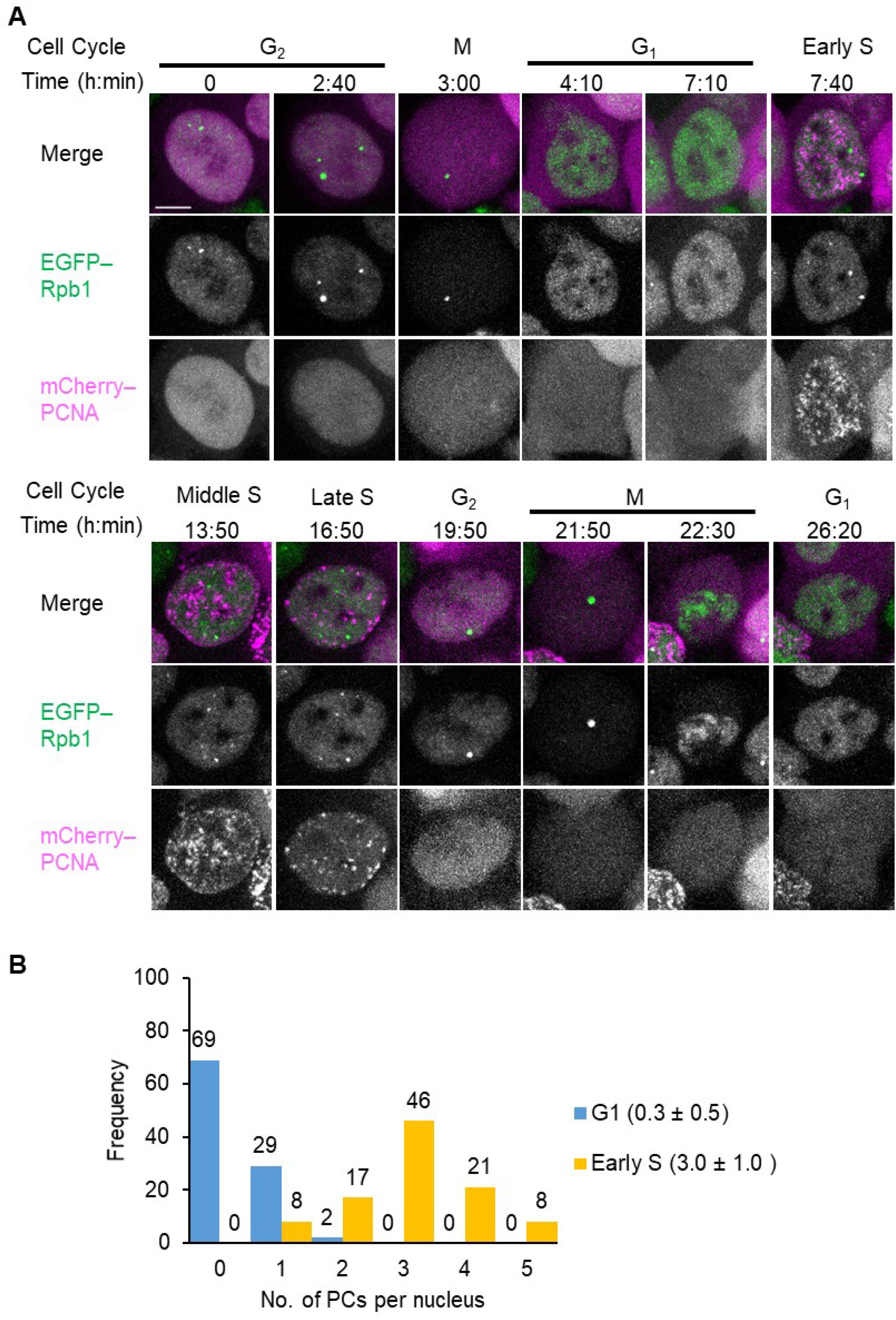
PC formation in living cells. **(A)** Time-lapse images of EGFP–Rpb1 and mCherry–PCNA in KI cells expressing mCherry– PCNA. MIP images of 15 Z-stacks at 0.80 μm intervals are shown. Scale bars, 5 μm. **(B)** Histogram (left) and average ±SD (right, n = 100 cells) of the number of PCs per nucleus in KI cells in both G_1_ and early S phases.

### 2.2. Live-cell imaging analysis of the association between PCs, CBs, and HLBs

The association between PCs, CBs, and HLBs was examined by expressing C-terminally SNAP-tagged coilin (coilin–SNAP) in coilin-KO cells that were generated from KI cells and C-terminally HaloTag-tagged NPAT (NPAT–HaloTag) in KI cells (Figs. 1A and S3A– D). Time-lapse imaging (Movie S2) revealed that some PCs nucleated near CBs (∼38% of PCs in 75 cells, n = 232, Table S2; Fig. 3A, arrowhead), whereas the others appeared to be distant from CBs (∼62%; Fig. 3A, asterisk). In some rare cases (∼3%), small PCs that nucleated away from CBs became bigger after associating with CBs (Fig. 3A, arrow). In NPAT–HaloTag expressing KI cells, PCs also nucleated near HLBs, but at a higher rate (∼51% PCs in 75 cells, n = 226, Table S2; Fig. 3B, arrowhead; and Movie S3). Some PCs simultaneously formed with HLBs (∼25%, Fig. 3B, arrow; and Movie S4) while others nucleated away from HLBs (∼24%; Fig. 3B, asterisk). These observations indicated that PCs preferentially nucleate near or together with HLBs.

**Figure 3.**
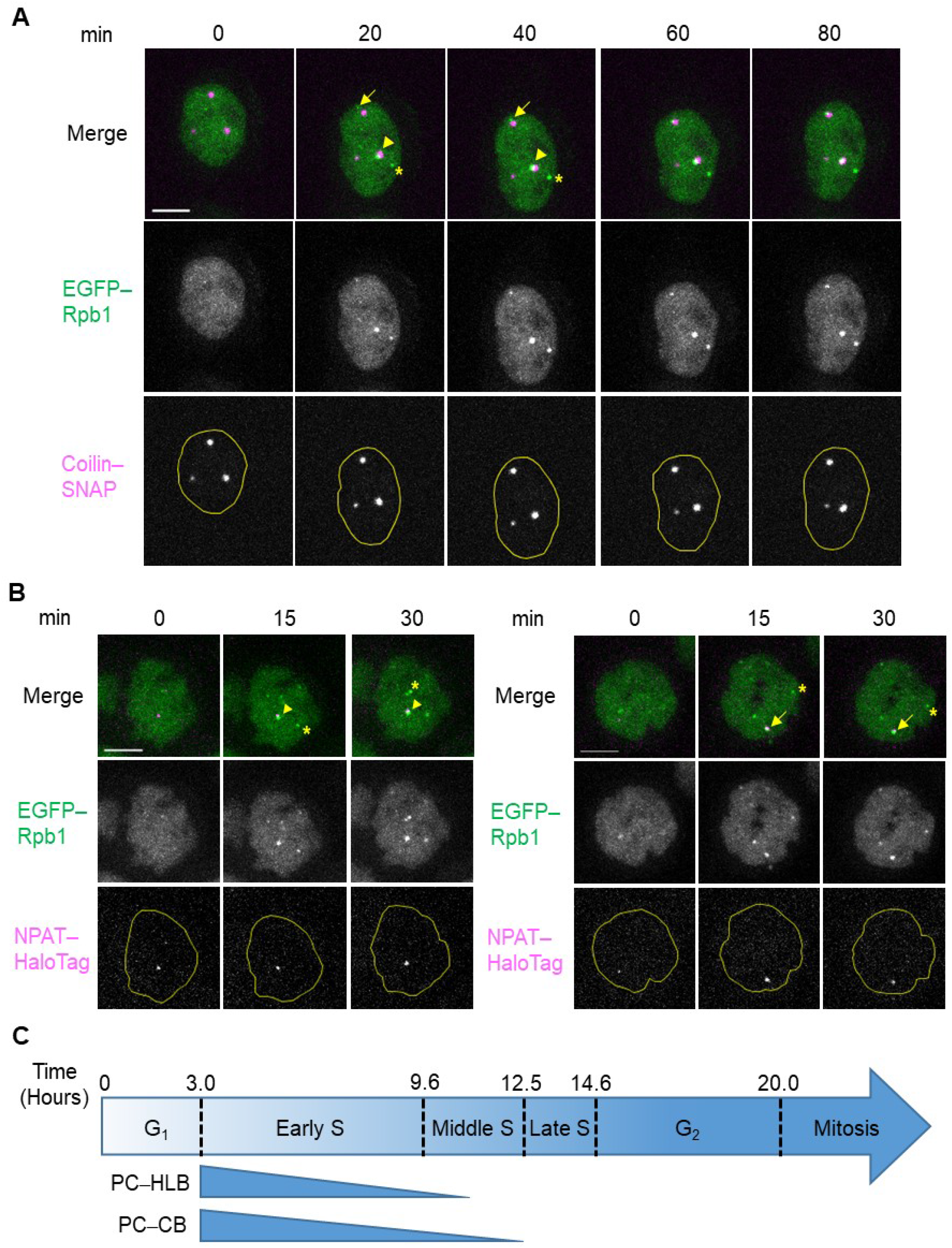
PCs nucleated near CBs and HLBs. **(A)** Time-lapse images of coilin-KO cells expressing coilin–SNAP. Confocal sections for EGFP–Rpb1 (Pol II; green in merge) and coilin–SNAP (CBs) labeled with SNAP–Cell 647– SiR (magenta in merge) were acquired. MIP images of 17 Z-stacks at 0.90 μm intervals are shown. A PC nucleated near a CB (arrowhead) and another nucleated apart from a CB (asterisk). A small PC (arrow) became bigger after association with a CB. **(B)** Time-lapse images of KI cells expressing NPAT–HaloTag. Confocal sections for EGFP–Rpb1 (Pol II; green in merge) and NPAT–HaloTag (HLBs) labeled with Janelia Fluor 646 HaloTag ligand (magenta in merge) were acquired. MIP images of 15 Z-stacks at 0.75 μm intervals are shown. A PC (arrowhead) nucleated near an existing HLB and another (asterisk) nucleated apart from an HLB. A PC nucleated with an HLB simultaneously (arrow). Yellow lines indicate nuclear peripheries. Scale bars, 5 μm. **(C)** Schematic representation of the association between PCs and CBs or HLBs during the cell cycle. PCs nucleated at the entry of S phase (Fig. 2A). The majority of PC–CB associates (90%) and PC–HLB associates (76%) dissolved during early S phase. Residual fractions of PC–CB associates (10%) and PC–HLB associates (24%) dissolved by the end of middle S phase. See also Table S3.

We next analyzed how long the PC–CB and PC–HLB associates were maintained using the time-lapse data. The mean association periods were 4.4 ± 1.9 (range, 1.3–9.3) h for PC–CB (n = 29 associates in 22 cells) and 4.6 ± 2.7 (range, 1.0–9.0) h for PC–HLB (n = 33 associates in 22 cells) (Table S3). Most PC–CB (90%) and PC–HLB (76%) associates dissolved within ∼6.6 h of early S phase, as judged from the mCherry–PCNA pattern (Fig. 3C). The residual fractions (10% for PC–CB and 24% for PC–HLB) had dissolved by 9.5 h, before mCherry–PCNA exhibited late-replicating heterochromatic distribution (Fig. 3C). PC–CB associates showed four patterns of dissolution: disappearance of PCs in the presence of CBs (38%), disappearance of CBs in the presence of PCs (21%), simultaneous disappearance of both (34%), and separation of a PC–CB associate into two distinct NBs (7%; n = 29 associates in 22 cells, Table S3). PC–HLB associates showed three patterns of dissolution: disappearance of PCs (46%), disappearance of HLBs (12%), and disappearance of both (42%, n =33 associates in 22 cells; Table S3). These findings indicated that PCs were less stable than CBs or HLBs.

The relationship between the three NBs (PCs, CBs, and HLBs) was examined using live-cell imaging with coilin-KO cells that expressed both coilin–SNAP and NPAT–HaloTag (Fig. 1A). These cells showed similar numbers of CBs or HLBs per nucleus to those of parental KI cells (Table S1). PCs in living cells were categorized into four patterns (n = 382 PCs in 100 cells; Fig. 4A): not associating with CBs or HLBs (17%), associating with only CBs (11%), associating with only HLBs (46%), and associating with both CBs and HLBs (26%). This observation was consistent with the higher rate of PC nucleation nearer HLBs than CBs, as observed previously (Fig. 3A, B, and Table S2). PC–CB–HLB associates (n = 142 in 100 cells imaged for 40 h) were reverse-tracked to reveal how ternary NB associations were formed. There were three association pathways (Fig. 4, B and C, pathways A–C; and Movie S5–7). In more than a half of all cases (53%), PCs and HLBs were simultaneously nucleated near CBs (pathway A; Movie S5). The second major pathway (32%) involved a preformed PC–HLB becoming associated with a nearby CB (pathway B; Movie S6). Nucleation of a PC near a preformed CB–HLB associate (pathway C; Movie S7) was relatively less frequent (15%). These findings are also consistent with PCs preferentially associating with HLBs than with CBs, and suggest a role of CBs in enhancing PC and HLB formation (pathway A).

**Figure 4.**
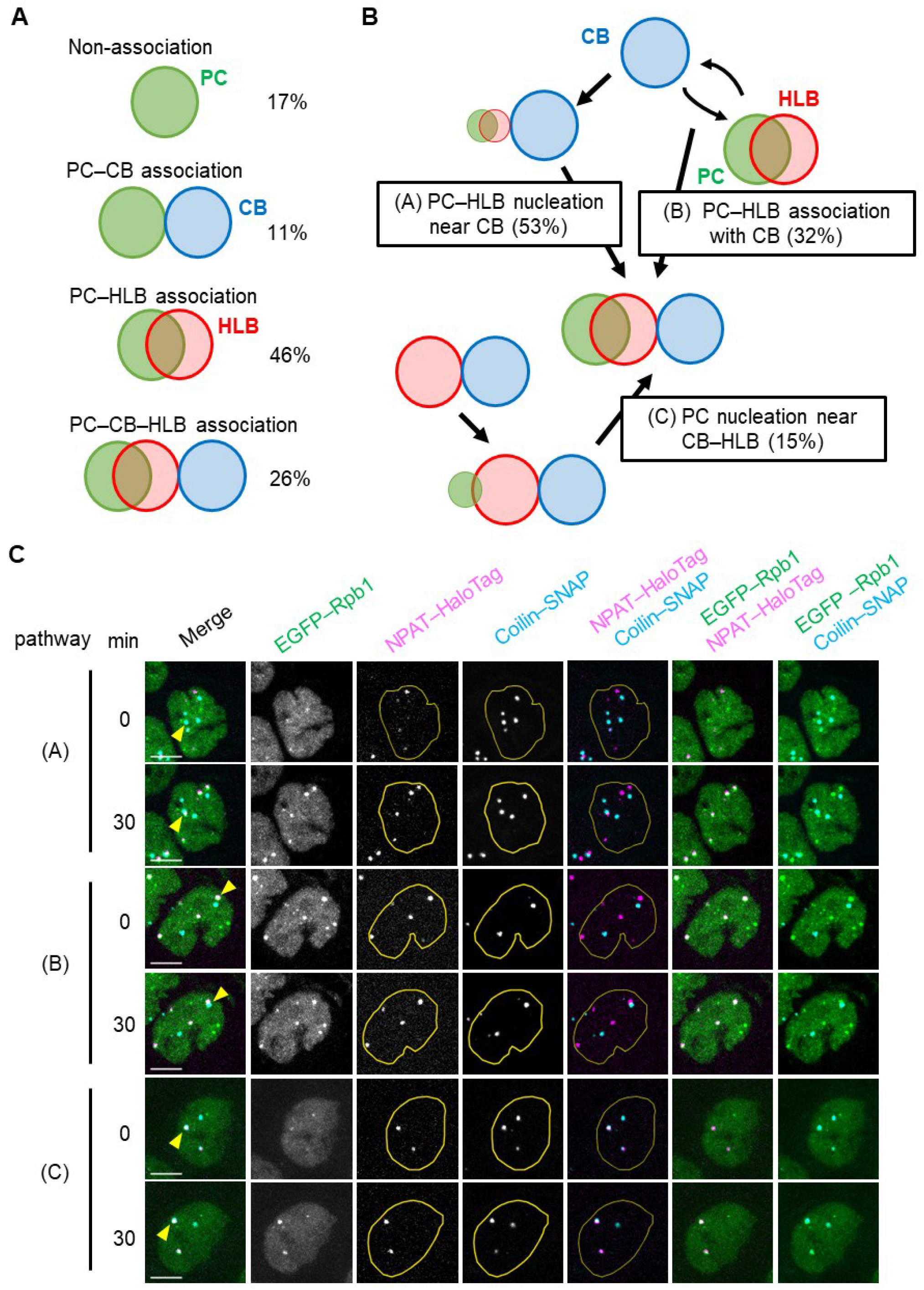
Association of NBs in living cells. **(A)** Association patterns between PCs, CBs, and HLBs, based on the analysis of 382 PCs in 100 coilin-KO cells expressing both NPAT–HaloTag and coilin–SNAP. **(B)** Schematic representation of three assembly pathways of PC–CB–HLB ternary associates. In this analysis, 142 PC–CB–HLB associates in 100 coilin-KO cells expressing both NPAT–HaloTag and coilin–SNAP were reverse-tracked from assembly of the ternary associates using live imaging data for 40 h. **(C)** Time-lapse images of EGFP–Rpb1 (green), NPAT–HaloTag (JF646; magenta), and coilin–SNAP (TMR; cyan) in coilin-KO cells expressing both NPAT–HaloTag and coilin– SNAP. MIP images of 13 Z-stacks at 0.75 μm intervals are shown. Yellow arrowheads indicate CBs associated with PCs and HLBs. Yellow lines indicate nuclear peripheries. Scale bars, 5 μm.

### 2.3. Coilin–KO did not affect PC formation but affected association between NBs

To investigate the requirement of CBs for PC formation, we used coilin-KO cells, which were generated from KI cells using CRISPR/Cas9 nickase (Figs. 1A and 5A, B). Coilin KO did not affect the number (5.4 ± 2.1 and 5.3 ± 1.9 in clones 1 and 2, respectively; Table S1) or phosphorylation status of PCs (Fig. S4A). Time-lapse imaging using coilin-KO cells that express mCherry–PCNA revealed that PCs appeared in early S phase (Fig. S5A and Movie S8), as observed in KI cells. These data indicate that CBs are not essential for PC formation. Similarly, HLBs and SMN foci were still observed in coilin-KO cells, and their numbers were unchanged compared with that of KI cells. By contrast, TCAB1 foci disappeared in coilin-KO cells (Fig. S5B and Table S1), supporting an essential role of CBs on TCAB1 foci [Chen et al, 2015; Vogan et al, 2016].

Next, by using immunofluorescence with fixed cells, we analyzed the associations between NBs in KI and coilin-KO cells in more detail. We classified the localization patterns of two protein pairs (i.e., Pol II–coilin, Pol II–NPAT, and coilin–NPAT) into three categories: “co-localizing” (most pixels overlapped), “adjacent” (few pixels overlapped), and “non-associating” (no overlapping pixels) (Fig. 5C). Both “co-localizing” and “adjacent” were treated as “associating,” which we used in live imaging as the spatial resolution was not high enough to distinguish “co-localizing” from “adjacent”. The co-localization rate between HLBs and PCs was higher than that between CBs and PCs in KI cells (Figs. 5D and S3D). In coilin-KO cells, HLBs and PCs became more co-localized, while the association between SMN foci and PCs decreased (Fig. 5D). These data suggest that the associations between PCs and NBs are balanced among different NBs, with a preferred association with HLBs, and so PC–HLB association becomes more prominent in the absence of CBs due to a lack of sequestration.

**Figure 5.**
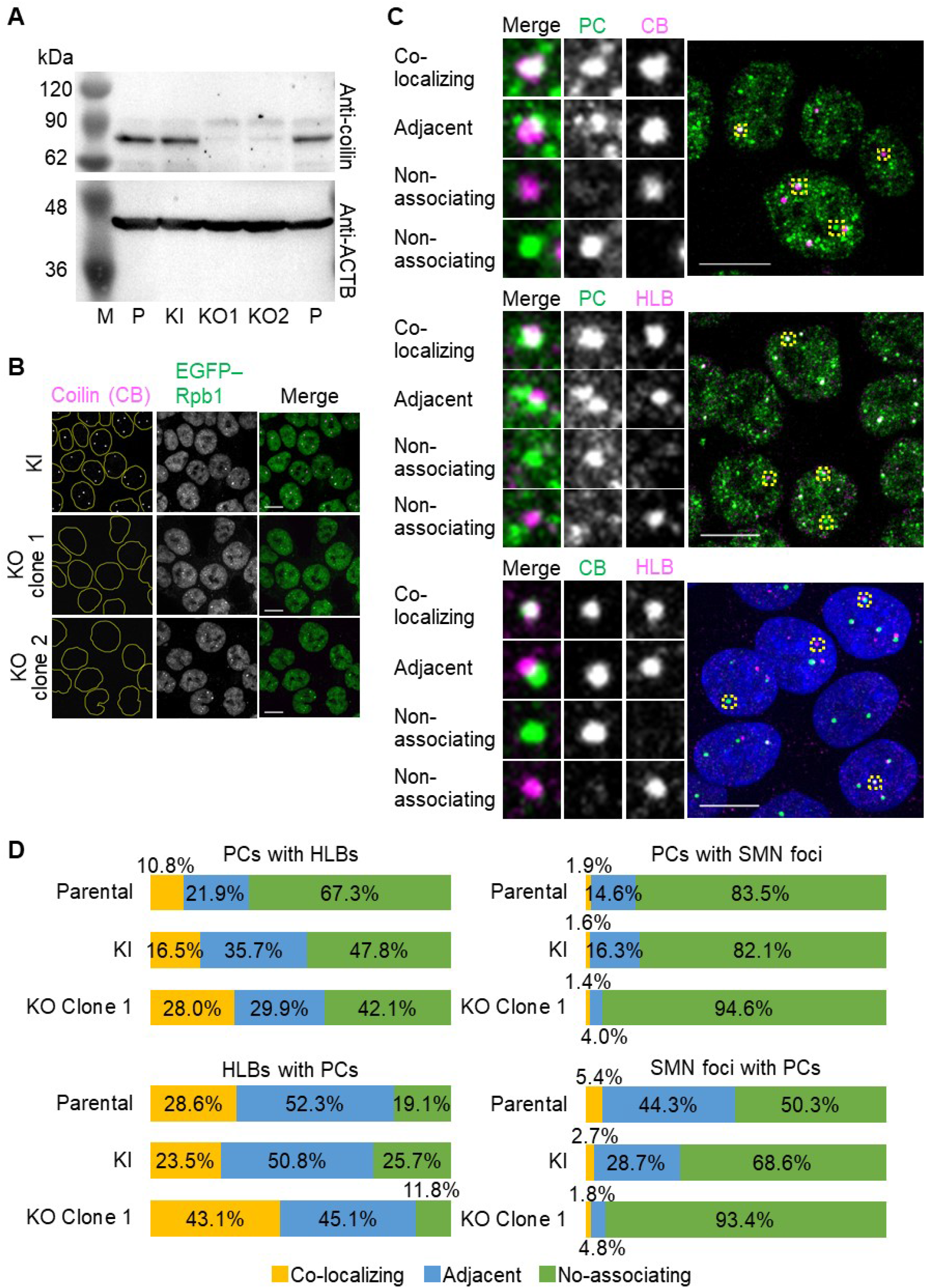
Effects of coilin–KO on PC formation and association with NBs. (**A**) and (**B**) Coilin-KO cells generated by CRISPR/Cas9 nickase system were analyzed by Western blotting **(A)** and immunofluorescence **(B). (A)** Whole-cell lysates separated on an SDS-polyacrylamide gel were probed with antibodies against coilin and ACTB. M, marker; P, parental HCT116 cells; KI, KI cells; KO1, coilin-KO cells clone 1; and KO2, coilin-KO cells clone 2. **(B)** Immunofluorescence images using antibodies against coilin in KI and coilin-KO cells. All images for coilin (magenta) and EGFP–Rpb1 (green) are MIP images of confocal Z-sections to cover the entire nucleus with 0.30 μm intervals (47–57 stacks). Yellow lines indicate nuclear peripheries. **(C)** and **(D)** Association of two heterotopic NBs. **(C)** Immunofluorescence. KI cells were fixed and stained with antibodies against coilin and NPAT to detect CBs and HLBs, respectively. PCs were detected using EGFP–Rpb1. Nuclear DNA was counterstained with Hoechst 33342 (only shown in cells stained with coilin and NPAT; blue). MIP images of Z-stacks at 0.30 μm intervals to cover the entire nucleus (51–58 stacks) are shown. The relationship between two different NBs was classified into three categories: overlapping, adjacent, and non-associating, and example images are shown. **(D)** Association rates between two heterotopic NBs. “PCs with HLBs or SMN foci” indicates that the percentage of PCs that were colocalized with, adjacent to, or non-associated with SMN foci or HLBs (top left and right). “HLBs or SMN foci with PCs” indicates the percentage of SMN foci or HLBs that were colocalized with, adjacent to, or non-associated with PCs (bottom left and right). Scale bars, 10 μm.

As CBs are known to co-localize with CDK7, which phosphorylates Ser5 in the Rpb1 CTD [Jordan et al, 1997], the association between PCs and CBs could facilitate Ser5 phosphorylation. However, levels of S5P in PCs with CBs were similar to those in PCs without CBs (Fig. S4B, C), suggesting that the function of the PC–CB association is not simply facilitating phosphorylation. In addition, levels of S5P in PCs were also constant, irrespective of the associations with HLBs (Fig. S4D). Thus, CBs and HLBs do not appear to play an essential role in regulating Ser5 phosphorylation in PCs.

### 2.4. PC depletion did not influence CBs and HLBs

We next investigated the effect of Pol II depletion on CBs and HLBs in parental HCT116 cells using triptolide, which degrades Pol II via Rpb1 ubiquitination [Wang et al, 2011]. When cells were treated with 5 μM triptolide for 2 h, Pol II signals detected using anti-CTD antibody were substantially diminished. In contrast, CBs and HLBs remained present in triptolide-treated cells (Fig. S6A), and the numbers of CBs and HLBs (average 3.3 and 2.3, respectively) were similar to those with dimethyl sulfoxide (DMSO)-treated control (average 3.4 and 2.2, respectively). The association between CBs and HLBs was also unaffected (Fig. S6B). Thus, the integrity of CBs and HLBs was independent of Pol II, consistent with the frequent nucleation of PCs from these NBs, but not vice versa.

### 2.5. PCs were liquid-like NBs compared with solid-like CBs and HLBs

Phase-separated NBs exhibit a broad range of biophysical properties, from solid-like to liquid-like [Boeynaems et al, 2018]; therefore, we investigated the sensitivity of PCs, CBs, and HLBs to 1,6-HD, which is known to disrupt liquid-like droplets rather than solid-like materials [Kroschwald et al, 2017]. PCs disappeared within 1 min after administration of 1,6-HD, whereas CBs and HLBs remained unchanged in coilin-KO cells expressing coilin–SNAP and KI cells expressing NPAT–HaloTag, respectively (Fig. 6), suggesting that PCs had more liquid-like properties compared with CBs and HLBs.

**Figure 6.**
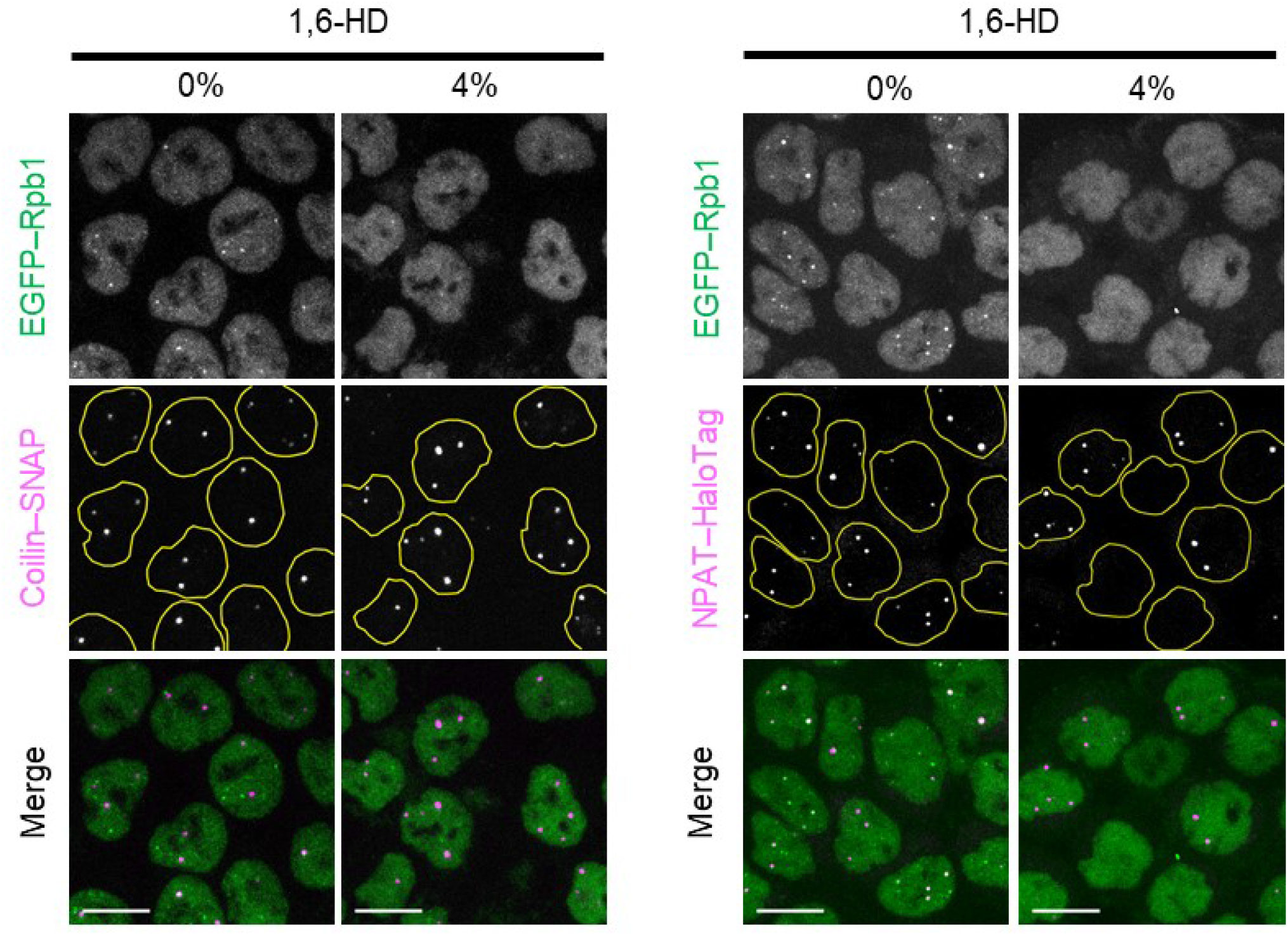
1,6-HD only disrupted PCs in living cells. Confocal images of coilin-KO cells expressing coilin–SNAP (left) and KI cells expressing NPAT–HaloTag (right) before and 1 min after the administration of 1,6-HD. MIP images of 11 Z-stacks at 0.90 μm intervals are shown for EGFP–Rpb1 (Pol II; green in merge) and coilin– SNAP (CBs) labeled with SNAP–Cell 647–SiR (magenta in merge) or NPAT–HaloTag (HLBs) labeled with Janelia Fluor 646 HaloTag ligand (magenta in merge). Yellow lines indicate nuclear peripheries. Scale bars, 10 μm.

## 3. DISCUSSION

The present study analyzed the formation of PCs, CBs, and HLBs, and their associations in HCT116 cells. PCs were often nucleated at the beginning of S phase near CBs and HLBs, and PC–CB and PC–HLB associates were dissolved before the end of S phase (Figs. 2 and 3; Tables S2 and S3). This observation is consistent with the function of CBs and HLBs, which are localized near sn/snoRNA and/or histone gene clusters and are involved in transcriptional and post-transcriptional regulation of these genes [Wang et al, 2016]. CBs were reported to associate with *HIST2* more frequently during late G_1_ and S phase compared with other cell cycle phases [Shopland et al, 2001], and HLBs became apparent during S phase [Duronio and Marzluff, 2017]. CBs and HLBs can be formed by increased local concentrations of the component proteins that bind to specific DNA sequence and/or RNA elements at these gene clusters. Once CBs and HLBs are formed, they are quite stably maintained probably due to their rather solid-like droplet nature, relatively resistant to 1,6-HD, through pi/pi, cation/pi, and electrostatic interactions [Ghule et al, 2008; Kaiser et al, 2008; Carmo-Fonseca and Rino, 2011; Shevtsov and Dundr, 2011; Alberti et al, 2019].

CBs and HLBs appear to facilitate the nucleation of PCs near the target gene cluster, such as sn/snoRNA genes (*RNU2* and *SNORD3A*) on chromosome 17 for CBs and the histone gene cluster, *HIST1*, on chromosome 6 for HLBs. Taken together with the fluorescence in situ hybridization data [Wang et al, 2016], the ternary associate of CBs, HLBs, and PCs is likely to be formed near chromosome 1 on which both sn/snoRNA genes and histone genes (*RNU1, RNU11, SNORA72, HIST2*, and *H3F3A*) are located (Fig. 7). A preformed CB on the sn/snoRNA gene locus can act as a core to form a PC–CB–HLB ternary associate via interaction with the histone gene locus, either by stimulating HLB and PC nucleation (Fig. 4B, C; pathway A) or interacting with a PC–HLB associate (Fig. 4B, C; pathway B). The presence of Pol II harboring S5P and S7P on its CTD in PCs also supports the view that PCs may contribute to the active transcription of sn/snoRNA and histone genes because these phosphorylated forms of Pol II (but not S2P) transcribe these genes (Fig. 1D) [Medlin et al, 2005; Egloff et al, 2007; Egloff et al, 2012]. Although CBs are not essential for the transcription of snRNA and histone genes, as coilin-KO cells grow normally [Tucker et al, 2001; Lemm et al, 2006; Whittom et al, 2008], CBs may contribute to the fine-tuning of sn/snoRNA transcription and the formation of other NBs, which may be required for precise developmental regulation [Tucker et al, 2001; Walker et al, 2009; Strzelecka et al, 2010].

**Figure 7.**
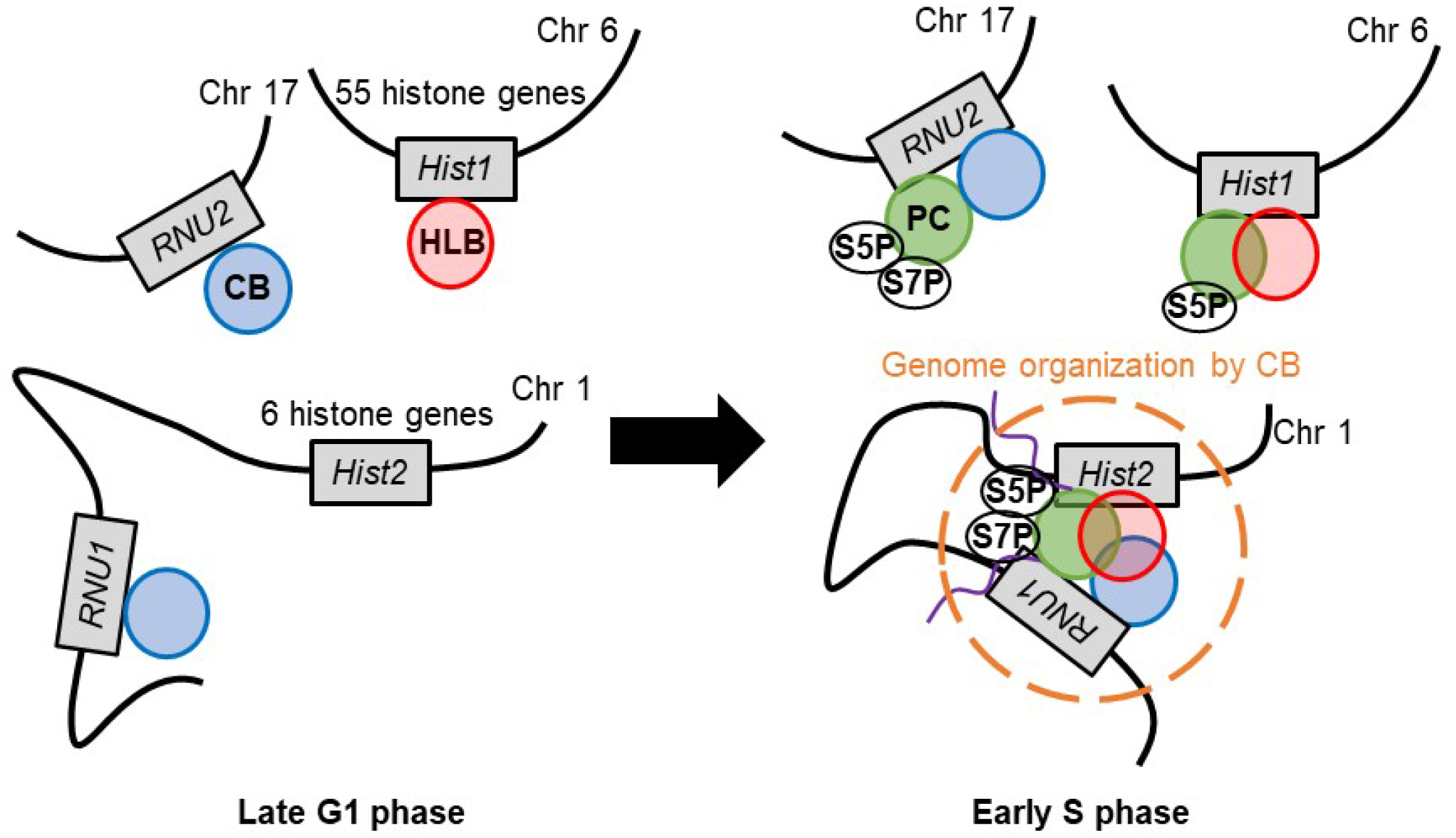
The model of the spatiotemporal and functional relationships between PCs, CBs, and HLBs. CBs and HLBs can be formed near sn/snoRNA and histone gene clusters with or without PCs. When histone genes are activated in S phase, the number of PCs increases in association with HLBs. CBs can organize histone genes and sn/snoRNA genes on chromosome 1 by forming ternary CB–HLB–PC associates. This genome organization may fine-tune transcription efficiency by Pol II molecules with S5P or S7P CTD in PCs.

Disruption of CBs via coilin–KO did not affect PCs in terms of the number per nucleus, nucleation timing, and the phosphorylation status of Pol II (Figs. S4 and S5; Table S1; Movie S8). However, the association between PCs and HLBs increased in response to coilin KO (Fig. 5D). This may be explained by a loss of PC sequestration to CBs. In the absence of CBs, increased free Pol II could facilitate nucleation at HLBs, and/or PCs free of CBs could associate with HLBs. Although it was of interest to investigate the effect of HLB depletion on PCs and CBs, this experiment was not possible in the present study since NPAT is essential for cell cycle progression [Di Fruscio et al, 1997; Ma et al, 2000; Ye et al, 2003]. In contrast to the increased association between PCs and HLBs, the association between PCs and SMN foci decreased by coilin KO (Fig. 5D), suggesting that a CB mediates the interaction between a SMN focus and a PC. Therefore, the presence or absence of one type of NB may affect other NBs via direct or indirect associations between NBs. A specific interaction between proteins in different NBs could mediate these associations. However, recent studies have suggested that interactions between different NBs may depend more on their biophysical properties than specific protein-protein interactions. Relative surface tensions are thought to be a key parameter in regulating associations between two heterotopic NBs [Bergeron-Sandoval et al, 2016; Ferric et al, 2016; Sawyer et al, 2016; Shin and Brangwynne, 2017]. Alterations of relative surface tensions among CBs, HLBs, and PCs may result in different association patterns. Although we observed overlapping and adjacent associations between NBs, the biological significance of these dynamic interactions remains unclear. Biological surfactant proteins may modulate the interactions between these NBs [Cuylen et al, 2016; Ferric et al, 2016].

## 4. EXPERIMENTAL PROCEDURES

### 4.1. Cell culture, transfection, and flow cytometry

HCT116 cells were obtained from ATCC (CCL-247, ATCC, VA, USA) and grown in high-glucose Dulbecco’s Modified Eagle Medium (DMEM; 08458-16, NACALAI TESQUE, Kyoto, Japan) supplemented with 10% fetal bovine serum (10270-106, Thermo Fisher Scientific, MA, USA), 2 mM L-glutamate, 100 U/mL penicillin, and 100 μg/mL streptomycin (G6784-100ML, Merck, Sigma-Aldrich, MO, USA) under 5% CO_2_ at 37°C. Lipofectamine 3000 was used for transfection according to the manufacturer’s instructions (L3000015, Thermo Fisher Scientific, MA, USA). Antibiotics selection was performed by culturing cells in medium supplemented with 500 μg/mL G418 for one week or 2 μg/mL puromycin for two days. A flow cytometer (SH800S, Sony, Tokyo, Japan) was used to collect cells expressing fluorescence-tagged proteins.

### 4.2 Plasmid construction, genome editing, and single-cell cloning

Total RNAs were extracted for use as cDNA synthesis templates using TRIzol (15596026, Thermo Fisher Scientific, MA, USA), treated with RQ1 RNase-Free DNase (M6101, Promega, WI, USA) in the presence of RNaseOUT (10777019, Thermo Fisher Scientific, MA, USA), extracted using phenol/chloroform, and precipitated using ethanol according to the manufacturers’ instructions. Primers to amplify cDNA were designed using Primer-BLAST (https://www.ncbi.nlm.nih.gov/tools/primer-blast/index.cgi). cDNA was synthesized using a PrimeScript II High Fidelity One Step RT-PCR Kit (R026A, Takara Bio, Shiga, Japan), purified using a QIAquick PCR Purification Kit (QIAGEN, 28106), and cloned into the PiggyBac Transposon Vector (PB533A-2, Funakoshi, Tokyo, Japan) or its derivatives with appropriate antibiotics resistant genes (ampicillin for *Escherichia coli* and neomycin or puromycin for mammalian cells) and a tag (SNAP-tag or HaloTag) using an In-Fusion HD Cloning Kit (639635, Takara Bio, Shiga, Japan). The *E. coli* DH5α strain was transformed using In-Fusion products, and single colonies were picked up for plasmid preparation and verification by DNA sequencing. For transfection, plasmids were purified using a Plasmid DNA Midiprep Kit (K210005, Thermo Fisher Scientific, MA, USA).

KI cells were generated by using TALE nuclease (TALEN) knock-in method using a Platinum Gate TALEN Kit, a gift from Takashi Yamamoto (1000000043, Addgene, MA, USA) [Sanjana et al, 2012]. To construct a TALEN pair that targets the 1st exon of POLR2A, DNA-binding modules for 5’-ATGCACGGGGGT-3’ and 5’-GCATGCGCTGTC-3’ were assembled into the ptCMV–153/47 vector, respectively. To construct a donor vector (pDonor–EGFP– POLR2A), ∼1,000 bp upstream of the annotated start codon of POLR2A, EGFP with A206K mutation, ∼250 bp fragment of the 1st exon including coding sequence, loxP-puromycin selection marker-loxP, and ∼1,000 bp of downstream sequence were tandemly flanked by PCR and inserted into the pBlueScript vector. The TALEN plasmids and pDonor–EGFP–POLR2A were co-transfected into parental HCT116 cells by electroporation using an NEPA21 super electroporator (Nepa Gene, Chiba, Japan). Puromycin-resistant clones were isolated and validated by genomic PCR, DNA sequencing, and western blotting. Note that KI cells used in this study spontaneously lost the puromycin cassette at the loxP site for unknown reasons.

Coilin-KO cells were generated using the Cas9 nickase system (pX335 and pKN7; Addgene) [Cong et al, 2013] with single guide RNAs designed using CHOPCHOP (http://chopchop.cbu.uib.no/). KO of the coilin gene (*COIL*) was achieved using the following sequences: 5′-TGCCTCAGGTGCGCGGCGCA<u>GGG</u>-3′ (forward) and 5′-TCCGCGCGCGAGAGCCGCCC<u>CCC</u>-3′ (reverse), where the PAM sequence is underlined. For single-cell cloning, transfected cells were resuspended in DMEM, filtered through a 40 μm cell strainer (352340, Corning), and seeded at a density of 1,000 cells/20 mL in a 15 cm dish. After incubation in a CO_2_ incubator for a week, single colonies were picked using a P20 pipette tip under a conventional phase–contrast microscope and transferred to an ibidi-coated 96-well plate (µ-Plate 96 Well Black, 89626, ibidi).

### 4.3. Antibodies, dyes, florescent ligands, and chemicals

All antibodies, dyes, and florescence ligands are listed in Table S4. Three mouse α-Rpb1 CTD monoclonal antibodies (IgGs) were generated in our laboratory [Stasevich et al, 2014]. The mouse monoclonal α-CTD antibody (C13B9) was generated using the unphosphorylated synthetic peptide corresponding to the CTD heptarepeat; however, the antibody can also recognize phosphorylated CTD [Stasevich et al, 2014]. Fab preparation and dye-conjugation were performed as previously described [Hayashi-Takanaka et al, 2011; Stasevich et al, 2014]. 1,6-HD was purchased from Tokyo Chemical Industry (H0099, Tokyo, Japan), dissolved in DMEM, filtered through a 0.2 μm filter for sterilization, and stored at 4°C. Triptolide was purchased from Tocris Bioscience (3253, Tocris, R&D Systems, MN, USA), dissolved in DMSO at 100 μM, and stored at −30°C. Triptolide was diluted in cell culture medium immediately prior to use.

### 4.4. Immunofluorescence, live-cell imaging, and image analysis

All immunofluorescence procedures were performed at room temperature. Cells were fixed in 4% paraformaldehyde, 0.1% Triton X-100, and 250 mM HEPES, pH 7.4 for 20 min, rinsed three times in Dulbecco’s phosphate-buffered saline (D-PBS) (−) (048-29805, FUJIFILM Wako Pure Chemical, Osaka, Japan), and permeabilized with 1% Triton X-100 in D-PBS (−) for 20 min. After blocked with Blocking One P (05999-84, NACALAI TESQUE, Kyoto, Japan) for 20 min and rinsed in D-PBS (−), the processed cells were incubated with primary antibodies in D-PBS (−) for 2 h. Cells were rinsed three times in D-PBS (−), incubated with secondary antibodies and Hoechst 33342 (H3570, Thermo Fisher Scientific) in D-PBS (−) for 2 h, and rinsed three times in D-PBS (−).

For live-cell imaging, cells were seeded on ibidi-treated culture dishes (µ-Slide 8 Well, 80826, or µ-Dish 35 mm high, 81156, ibidi) and cultured for one to two days. The medium was replaced with FluoroBright (A1896701, Thermo Fisher Scientific containing 10% fetal bovine serum, 2 mM L-glutamate, 100 U/mL penicillin, and 100 μg/mL streptomycin, immediately prior to imaging. Images were acquired using a spinning disc confocal microscope comprising an inverted microscope (Ti-E, Nikon), a Plan-Apo VC 100x (NA 1.4) oil immersion objective lens (NA 1.4; with Type F immersion oil, MXA22168, Nikon), a spinning disk unit (CSU-W1, Yokogawa), an EM-CCD camera (iXon3 DU888 X-8465, Andor), and a LU-N4 laser unit (Nikon) under the control of an operating software (NIS-Elements, Nikon). Live-cell imaging was performed at 37°C under 5% of CO_2_ using a stage top incubator (INUBG2TF-WSKM-A14R, Tokai-Hit). Images were analyzed using Icy [Chaumont et al, 2012] (http://icy.bioimageanalysis.org/). Individual Z-section images were used for quantitative analysis, although Max-Intensity-Projection (MIP) images of Z-series were presented in all figures unless stated otherwise. To detect foci of NBs, the threshold was set as 30 percentiles intensity to cut off pixels with intensity <30% of an intensity histogram of all pixels in an image with a size filer between 10 and 150 pixels in order. All results were checked visually and, in cases in which 30 percentiles did not sufficiently detect foci, the threshold value was changed manually. The distance threshold judged as “co-localizing” was within 3 pixels (390 nm) between centers of foci in 3D. We analyzed cells that exhibited at least one PC, CB, HLB, SMN focus, or TCAB1 focus.

### 4.5. Western blotting

Cells were trypsinized, collected into 15 mL tubes, and washed twice in 10 mL PBS (−). The cell density was counted and lysed in Laemmli buffer without dithiothreitol and bromophenol blue to yield 1 × 10^7^ cells/mL. Whole-cell lysates were heated at 95°C for 10 min and stored at −30°C. Protein concentration was measured using a bicinchoninic acid assay kit (297-73101, FUJIFILM Wako Pure Chemical, Osaka, Japan). After supplemented with dithiothreitol and bromophenol blue, and boiling at 95°C for 5 min, whole lysates (50 μg to detect coilin or 90 μg to detect Rpb1 and GFP) were separated by sodium dodecyl sulfate (SDS)-polyacrylamide gel electrophoresis (90 min at 20 mA constant currency per 85 mm height gel). Gels were gently shaken in Transfer buffer (100 mM Tris, 200 mM glycine, and 5% methanol) for 10 min. Polyvinylidene fluoride membranes (BSP0161, FluoroTrans W PVDF Transfer Membranes, Pall, NY, USA) were immersed in methanol for 3 min and equilibrated in Transfer buffer for 10 min with gentle shaking. Proteins on gels were transferred to membranes for 60 min at 150 mA (for coilin) or 250 mA (for Rpb1) per 85 × 90 mm membrane. After an incubation in Blocking One (03953-95, NACALAI TESQUE, Kyoto, Japan) for 20 min with gentle shaking, membranes were incubated with primary antibodies diluted in Can Get Signal Solution I (NKB-101, TOYOBO, Osaka, Japan) overnight at 4°C (for coilin and GFP) or 2 h at room temperature (for Rpb1). After washed in TBS-T (100 mM Tris-HCl, pH 8.0, 0.1% Tween-20, 150 mM NaCl) for 20 min three times, membranes were incubated with horseradish peroxide-conjugated secondary antibodies (80 ng/mL, AB_10015289, Jackson ImmunoResearch, PA) in Can Get Signal Solution II (NKB-101, TOYOBO, Osaka, Japan) for 1 h at room temperature. After membranes were washed in TBS-T for 20 min three times, signals were developed using ImmunoStar LD (296-69901, FUJIFILM Wako Pure Chemical, Osaka, Japan) and detected using a gel documentation system (LuminoGraph II, ATTO).

## ACKNOWLEDGEMENTS

We thank Cristina Cardoso and Takashi Yamamoto for plasmid constructs. We also thank the Biomaterials Analysis Division, Open Facility Center, Tokyo Institute of Technology for DNA sequencing analysis. This work was in part of supported by MEXT/JSPS KAKENHI (JP17H01417, and JP18H05527 to H.K.; JP18H05498 to Y.S.), Japan Science and Technology Corporation (JST-CREST; JPMJCR16G1), and Japan Agency for Medical Research and Development (AMED-BINDS; JP20am0101105) to H.K. We are grateful to Yoshida Scholarship Foundation for financial support to T.I during his PhD.

## SUPPLEMENTARY MATERIALS

Figure S1. Phosphorylation status of Pol II CTD on PCs in parental HCT116 cells

Figure S2. NBs in parental HCT116 and KI cells

Figure S3. SNAP–coilin was more representative of the endogenous coilin than coilin–SNAP Figure S4. Phosphorylation status of Pol II CTD on PCs did not depend on CBs or HLBs

Figure S5. PCs in coilin-KO cells

Figure S6. Effects of PC depletion by triptolide treatment on CB and HLB association in parental HCT116 cells

Table S1. The number of NBs per nucleus in cells used in this study

Table S2. Nucleation patterns of PCs in coilin-KO cells expressing coilin–SNAP and in KI cells expressing NPAT–HaloTag

Table S3. Time course of dissolution of PC–CB associates and PC–HLB associates

Table S4. The list of antibodies and dyes used in this study

Movie S1. The time-lapse movie of KI cells expressing mCherry–PCNA

Movie S2. The time-lapse movie of coilin-KO cells expressing coilin–SNAP, showing a PC nucleated near a CB and another nucleated apart from a CB

Movie S3. The time-lapse movie of KI cells expressing NPAT–HaloTag, showing a PC nucleated near a preformed HLB

Movie S4. The time-lapse movie of KI cells expressing NPAT–HaloTag, showing a PC nucleated with an HLB simultaneously

Movie S5. The time-lapse movie of coilin-KO cells expressing both coilin–SNAP and NPAT– HaloTag, showing pathway A

Movie S6. The time-lapse movie of coilin-KO cells expressing both coilin–SNAP and NPAT– HaloTag, showing pathway B

Movie S7. The time-lapse movie of coilin-KO cells expressing both coilin–SNAP and NPAT– HaloTag, showing pathway C

Movie S8. The time-lapse movie of coilin-KO cells expressing mCherry–PCNA

## Legends to Supplementary Movies

**Movie S1. The time-lapse movie of KI cells expressing mCherry–PCNA**

Confocal images of EGFP–Rpb1 (green) and mCherry–PCNA (magenta) in KI cells expressing mCherry–PCNA were acquired using time-lapse microscopy (see also Fig. 2A). The movie is MIP of 15 Z-stacks at 0.80 μm intervals. Scale bar, 5 μm.

**Movie S2. The time-lapse movie of coilin-KO cells expressing coilin–SNAP, showing a PC nucleated near a CB and another nucleated apart from a CB**

Confocal images of EGFP–Rpb1 (green) and coilin–SNAP (magenta) in the cells were acquired using time-lapse microscopy (see also Fig. 3A). The movie is MIP of 17 Z-stacks at 0.90 μm intervals. Scale bar, 5 μm.

**Movie S3. The time-lapse movie of KI cells expressing NPAT–HaloTag, showing a PC nucleated near a preformed HLB**

Confocal images of EGFP–Rpb1 (green) and NPAT–HaloTag (magenta) in KI cells expressing NPAT–HaloTag were acquired using time-lapse microscopy (see also Fig. 3B left). The movie is MIP of 15 Z-stacks at 0.75 μm intervals. Scale bar, 5 μm.

**Movie S4. The time-lapse movie of KI cells expressing NPAT–HaloTag, showing a PC nucleated with an HLB simultaneously**

Confocal images of EGFP–Rpb1 (green) and NPAT–HaloTag (magenta) in KI cells expressing NPAT–HaloTag were acquired using time-lapse microscopy (see also Fig. 3B right). The movie is MIP of 15 Z-stacks at 0.75 μm intervals. Scale bar, 5 μm.

**Movie S5. The time-lapse movie of coilin-KO cells expressing both coilin–SNAP and NPAT–HaloTag, showing pathway A**

Confocal images of EGFP–Rpb1 (green), NPAT–HaloTag (JF646; magenta), and coilin–SNAP (TMR; cyan) in coilin-KO cells expressing both coilin–SNAP and NPAT–HaloTag. See also Fig. 4C, Pathway A. The movie is MIP of 13 Z-stacks at 0.75 μm intervals. Scale bar, 5 μm.

**Movie S6. The time-lapse movie of coilin-KO cells expressing both coilin–SNAP and NPAT–HaloTag, showing pathway B**

Confocal images of EGFP–Rpb1 (green), NPAT–HaloTag (JF646; magenta), and coilin–SNAP (TMR; cyan) in coilin-KO cells expressing both coilin–SNAP and NPAT–HaloTag. See also Fig. 4C, Pathway B. The movie is MIP of 13 Z-stacks at 0.75 μm intervals. Scale bar, 5 μm.

**Movie S7. The time-lapse movie of coilin-KO cells expressing both coilin–SNAP and NPAT–HaloTag, showing pathway C**

Confocal images of EGFP–Rpb1 (green), NPAT–HaloTag (JF646; magenta), and coilin–SNAP (TMR; cyan) in coilin-KO cells expressing both coilin–SNAP and NPAT–HaloTag. See also Fig. 4C, Pathway C. The movie is MIP of 13 Z-stacks at 0.75 μm intervals. Scale bar, 5 μm.

**Movie S8. The time-lapse movie of coilin-KO cells expressing mCherry–PCNA**

Confocal images of EGFP–Rpb1 (green) and mCherry–PCNA (magenta) in coilin-KO cells expressing mCherry–PCNA were acquired using time-lapse microscopy (see also Fig. S5A). The movie is MIP of 15 Z-stacks at 0.75 μm intervals. Scale bar, 5 μm.

**Table S1.** The number of NBs per nucleus in cells used in this study

| Cell line | PCs | CBs | HLBs | SMN foci | TCAB1 foci |
| --- | --- | --- | --- | --- | --- |
| Parental HCT116 | 8.9 ± 3.7 | 2.8 ± 1.2 | 3.5 ± 1.3 | 3.3 ± 1.9 | 3.4 ± 1.3 |
| KI | 5.4±2.2 | 2.8 ± 1.3 | 3.3 ± 1.1 | 2.6 ± 1.3 | 2.6 ± 0.9 |
| Coilin-KO | 5.4 ± 2.1 (Clone 1)<br>5.3 ± 1.9 (Clone 2) | N.D.<sup>a</sup> | 3.1 ± 1.0 (Clone 1) | 4.6 ± 3.2 (Clone 1) | N.D. |
| Coilin-KO expressing<br>SNAP–coilin | 5.0 ± 1.8 | 2.8 ± 1.9 | 3.3 ± 1.0 | N.D. | N.D. |
| Coilin-KO expressing<br>coilin–SNAP | 4.6 ± 1.7 | 3.6 ± 1.4 | 2.8 ± 1.1 | N.D. | N.D. |
| Coilin-KO expressing<br>coilin–SNAP and NPAT–HaloTag | N.D. | N.D. | 2.9 ± 0.9 | N.D. | N.D. |

<sup>a</sup>N.D., Not determined

**Table S2.** Nucleation patterns of PCs in coilin-KO cells expressing coilin–SNAP and in KI cells expressing NPAT–HaloTag.

| Coilin-KO cells expressing coilin–SNAP<br>(n = 232 PCs in 75 cells) | PCs nucleated near CBs | PCs nucleated away from CBs |  |
| --- | --- | --- | --- |
|  |  | PCs not associated throughout | PCs eventually associated with CBs |
|  | 38% | 59% | 3% |
| KI cells expressing NPAT–HaloTag<br>(n = 226 PCs in 75 cells) | PCs nucleated near HLBs | PCs nucleated with HLBs | PCs nucleated away from HLBs |
|  | 51% | 25% | 24% |

**Table S3.** Dissolution of NB associates

|  |  | PC–CB<sup>a</sup> | PC–HLB<sup>b</sup> |
| --- | --- | --- | --- |
| No. of NB associates (cells) |  | 29 (22) | 32 (22) |
| Dissolution time | by ~6.6 h (early S) | 90% | 76% |
|  | by ~9.5 h (middle S) | 10% | 24% |
| Dissolution pattern | PC disappeared | 38% | 46% |
|  | Both PC and CB–HLB disappeared | 34% | 42% |
|  | CB–HLB disappeared | 21% | 12% |
|  | PC–CB–HLB separated | 7% | 0% |

<sup>a</sup>Coilin-KO cells expressing coilin–SNAP
<sup>b</sup>KI cells expressing NPAT–HaloTag

**Table S4.** The list of antibodies and dyes used in this study

|  | Name | Clone | Supplier | Code | Immunofluorescence | Western Blotting |
| --- | --- | --- | --- | --- | --- | --- |
| Primary antibodies | Mouse anti-coilin | Pdelta | Santa Cruz Biotechnology | sc-56298 | 0.2 µg/mL | N.D.<sup>a</sup> |
|  | Mouse anti-coilin | F-7 | Santa Cruz Biotechnology | sc-55594 | N.D. | 1 µg/mL |
|  | Rabbit anti-coilin | N.D. | Proteintech | 10967-1-AP | 20,000x | N.D. |
|  | Mouse anti-fibrillarin | B-1 | Santa Cruz Biotechnology | sc-166001 | 0.3 µg/mL | N.D. |
|  | Mouse anti-SMN | 2B1 | Santa Cruz Biotechnology | sc-32313 | 0.2 µg/mL | N.D. |
|  | Mouse anti-NPAT | 27 | Santa Cruz Biotechnology | sc-136007 | 0.2 µg/mL | N.D. |
|  | Mouse anti-beta actin | AC-15 | Santa Cruz Biotechnology | sc-69879 | N.D. | 150 ng/mL for 2 h at R.T.<br>70 ng/mL for O/N at 4 deg. |
|  | Mouse anti-Rpb1 unphosphorylated CTD | C13B9 | Lab made | N.D. | 1 µg/mL | 1 µg/mL |
|  | Mouse anti-Rpb1 S2P CTD | Pc26 | Lab made | N.D. | 1 µg/mL | N.D. |
|  | Mouse anti-Rpb1 S5P CTD | Pa57 | Lab made | N.D. | 1 µg/mL | N.D. |
|  | Rat anti-Rpb1 S7P CTD | 4E12 | Active Motif | 61087 | 1 µg/mL | N.D. |
|  | Rat anti-Rpb1 T4P CTD | 6D7 | Active Motif | 61961 | 1 µg/mL | N.D. |
|  | Mouse anti-GFP | GF200 | Nacalai Tesque | 04363-24 | N.D. | 2 µg/mL |
| Secondary antibodies | Donkey anti-rabbit IgG |  | Jackson ImmunoResearch | 711-005-152 | 1 µg/mL | N.D. |
|  | Goat anti-rabbit IgG |  | Jackson ImmunoResearch | 111-005-144 | 1 µg/mL | N.D. |
|  | Goat anti-mouse IgG, Fc fragment specific |  | Jackson ImmunoResearch | 115-005-071 | 1 µg/mL | N.D. |
|  | HRP-conjugated goat anti-mouse IgG |  | Jackson ImmunoResearch | 115-035-003 | N.D. | 80 ng/mL |
|  | Donkey anti-rat IgG |  | Jackson ImmunoResearch | 712-005-153 | 1 µg/mL | N.D. |
|  | Donkey anti-mouse IgG |  | Jackson ImmunoResearch | 715-005-150 | 1 µg/mL | N.D. |
| Dyes | Alexa Fluoro 488 |  | Thermo Fisher Scientific | A30052 |  |  |
|  | Cy3 |  | GE Healthcare | Q13108 |  |  |
|  | Cy5 |  | GE Healthcare | Q15108 |  |  |
|  | Hoechst 33342 |  | Thermo Fisher Scientific | C10329 |  |  |
|  | SNAP–Cell TMR–Star |  | New England BioLabs | S9105 |  |  |
|  | SNAP–Cell 647–SiR |  | New England BioLabs | S9102 |  |  |
|  | Janelia Fluor 549 HaloTag |  | Promega | GA1111 |  |  |
|  | Janelia Fluor 646 HaloTag |  | Promega | GA1121 |  |  |

<sup>a</sup>N.D., Not done

**Figure S1.**
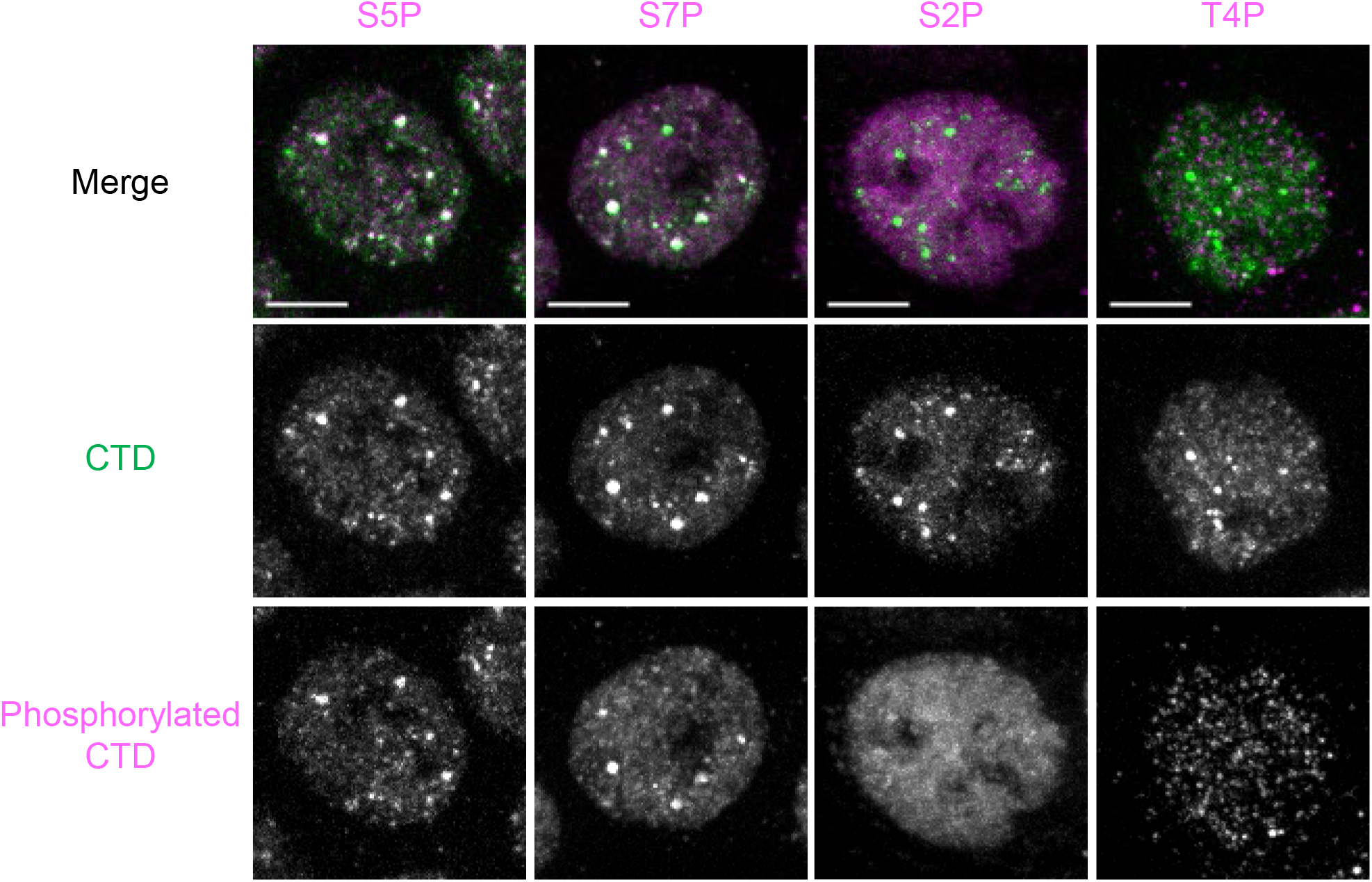
Phosphorylation status of Pol II CTD on PCs in parental HCT116 cells. Parental HCT116 cells were fixed and stained with antibodies against Pol II CTD (Alexa Fluor 488; green) and S2P, T4P, S5P, or S7P (Cy5; magenta). All images shown are MIP images of Z-stacks at 0.30 μm intervals to cover the entire nucleus (29–62 stacks). Scale bars, 5 μm.

**Figure S2.**
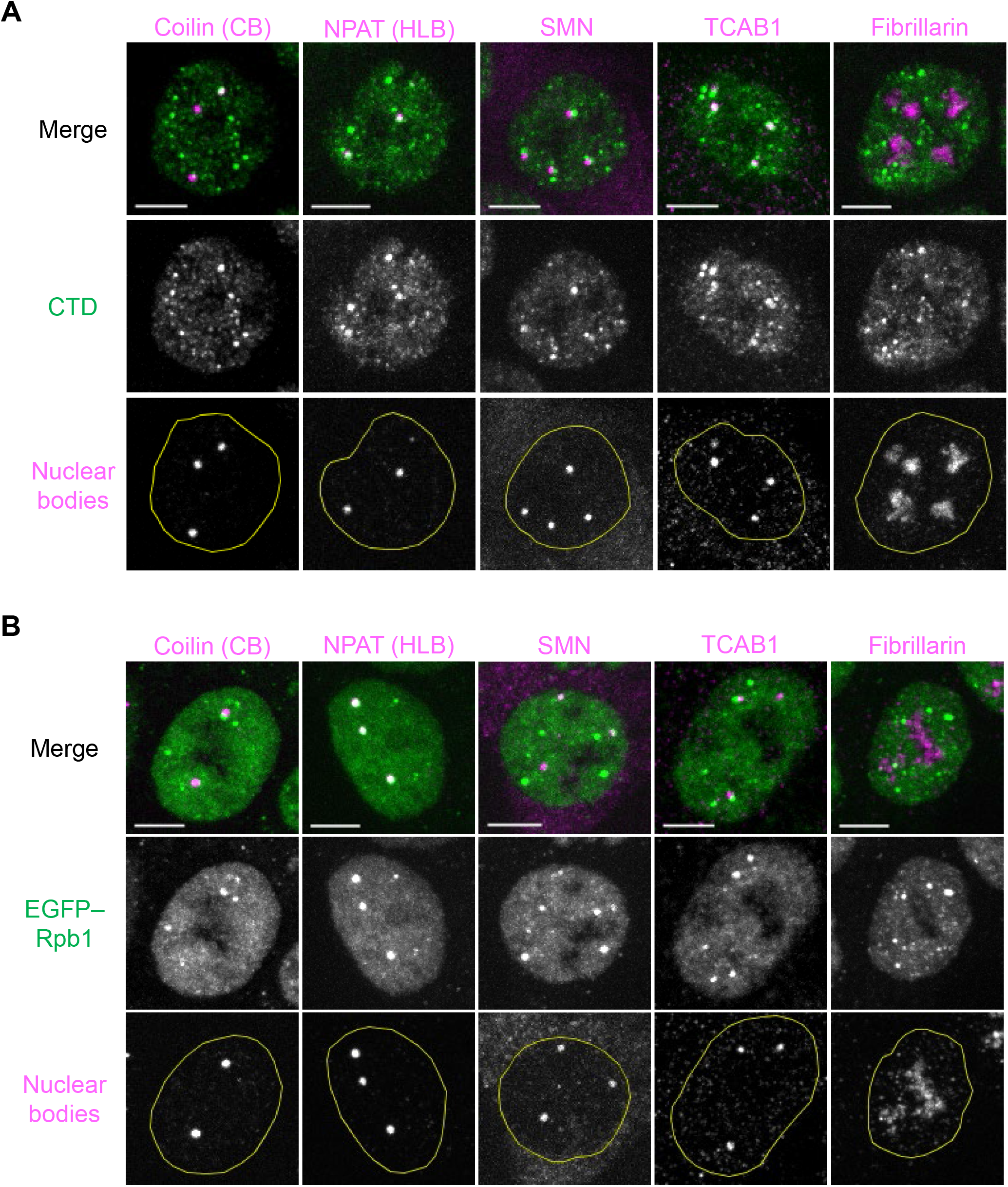
NBs in parental HCT116 and KI cells. Parental HCT116 and KI cells were fixed and stained with antibodies against coilin, SMN, TCAB1, fibrillarin, and NPAT, and Cy5-labeled secondary antibody (magenta). PCs were detected using Alexa Fluor 488-labeled anti-CTD in parental HCT116 cells **(A)** or with EGFP–Rpb1 in KI cells **(B)** (green). Confocal images of the entire nucleus were acquired at 0.30 μm Z-intervals (29–62 stacks). MIP images are shown. Yellow lines indicate nuclear peripheries. Scale bars, 5 μm.

**Figure S3.**
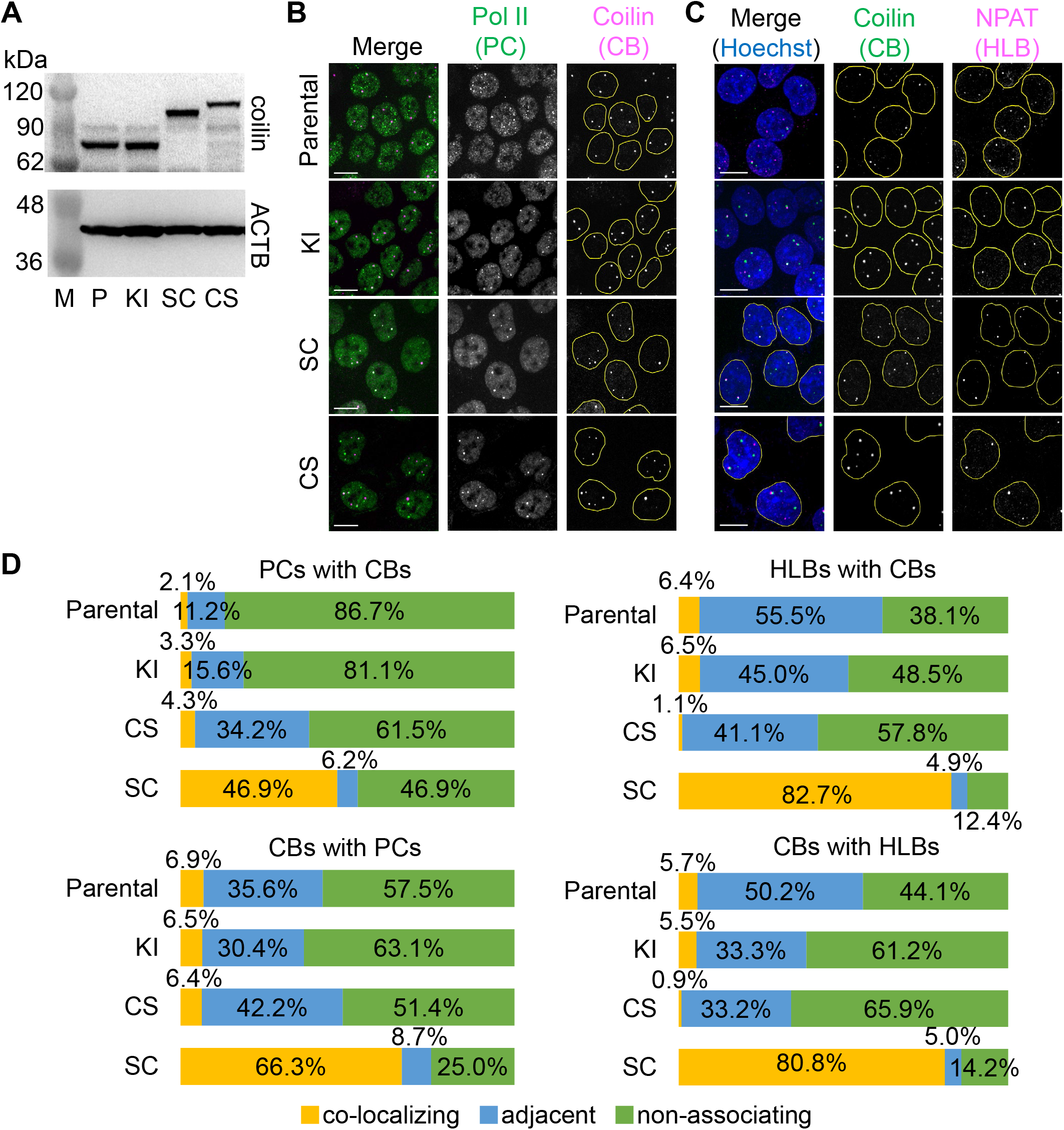
SNAP–coilin was more representative of the endogenous coilin than coilin–SNAP. Using KI cells, coilin-KO cells were first generated and then cells expressing SNAP-tagged coilin were established. N-terminally and C-terminally SNAP-tagged versions (SNAP–coilin and coilin–SNAP, respectively) were examined. **(A)** Western blotting. Whole-cell lysates were separated on an SDS-polyacrylamide gel and probed with antibodies against coilin and ACTB. M, marker; P, parental HCT116 cells; KI, KI cells; SC, coilin-KO cells expressing SNAP–coilin (clone 1); CS, coilin-KO cells expressing coilin–SNAP (clone 1). **(B)** and **(C)** Immunofluorescence. **(B)** Parental HCT116 cells were stained with antibodies against Pol II CTD (green) and coilin (magenta). EGFP–Rpb1 (green) in KI cells which were stained with antibodies against coilin (magenta). **(C)** Cells were stained with antibodies against coilin and NPAT to detect CBs (green) and HLBs (magenta), respectively. Nuclear DNA was counterstained with Hoechst 33342 (blue). Yellow lines indicate nuclear peripheries. Scale bars, 10 μm. **(D)** Association rates between two heterotopic NBs. “PCs or HLBs with CBs” indicates that the percentage of PCs or HLBs that were colocalized with, adjacent to, or non-associated with CBs (top left and right). “CBs with PCs or HLBs” indicates the percentage of CBs that were colocalized with, adjacent to, or non-associated with PCs (bottom left and right). Parental, parental HCT116 cells; KI, KI cells; SC, coilin-KO cells expressing SNAP–coilin (clone 1); CS, coilin-KO cells expressing coilin–SNAP (clone 1).

**Figure S4.**
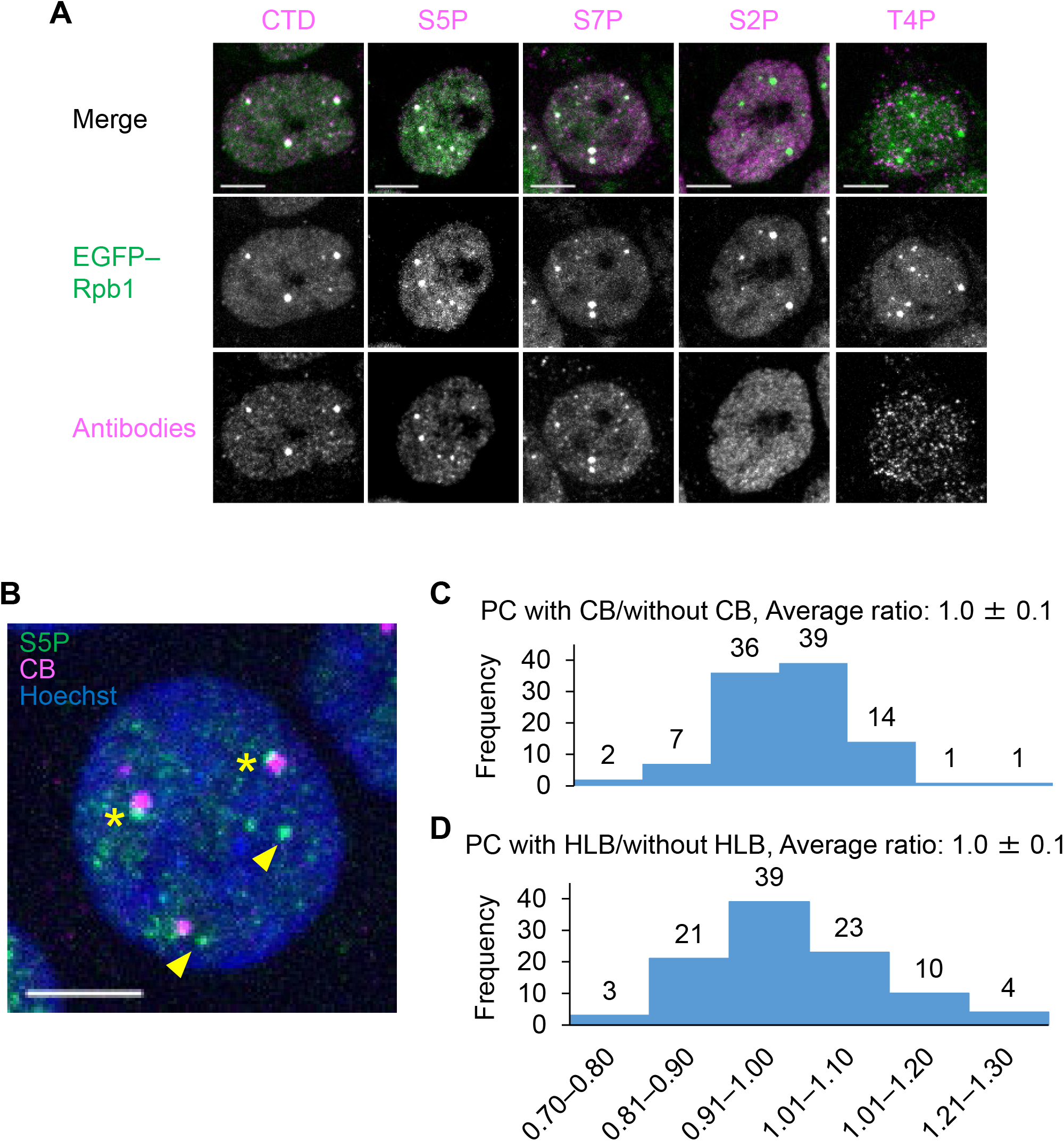
Phosphorylation status of Pol II CTD on PCs did not depend on CBs or HLBs. **(A)** Coilin-KO cells were fixed and stained with Pol II CTD antibodies (magenta). PCs detected using EGFP– Rpb1 (green) harbored S5P and S7P, as in coilin-positive KI cells. MIP images of Z-stacks at 0.30 μm intervals to cover the entire nucleus (29–62 stacks) are shown. **(B–D)** S5P levels in PCs with or without association with CBs or HLBs. **(B)** Immunofluorescence: Parental HCT116 cells were stained with antibodies against S5P (green) and coilin (magenta). Nuclear DNA was counterstained using Hoechst 33342 (blue). Arrowheads and asterisks indicate S5P foci (PCs) with and without a CB, respectively. An MIP image of 47 Z-stacks at 0.30 μm intervals is shown. **(C)** S5P levels in S5P foci with and without CBs. Intensity ratio of CB-associating PCs to CB-free PCs was expressed as “average intensity of a S5P focus with a CB in a cell” divided by “average intensity of a S5P focus without a CB in the same cell” (n = 100 cells). **(D)** S5P levels in S5P foci with and without HLBs. Intensity ratio of HLB-associating PCs to HLB-free PCs was expressed as “average intensity of a S5P focus with an HLB in a cell” divided by “average intensity of a S5P focus without an HLB in the same cell” (n = 100 cells). Scale bars, 5 μm.

**Figure S5.**
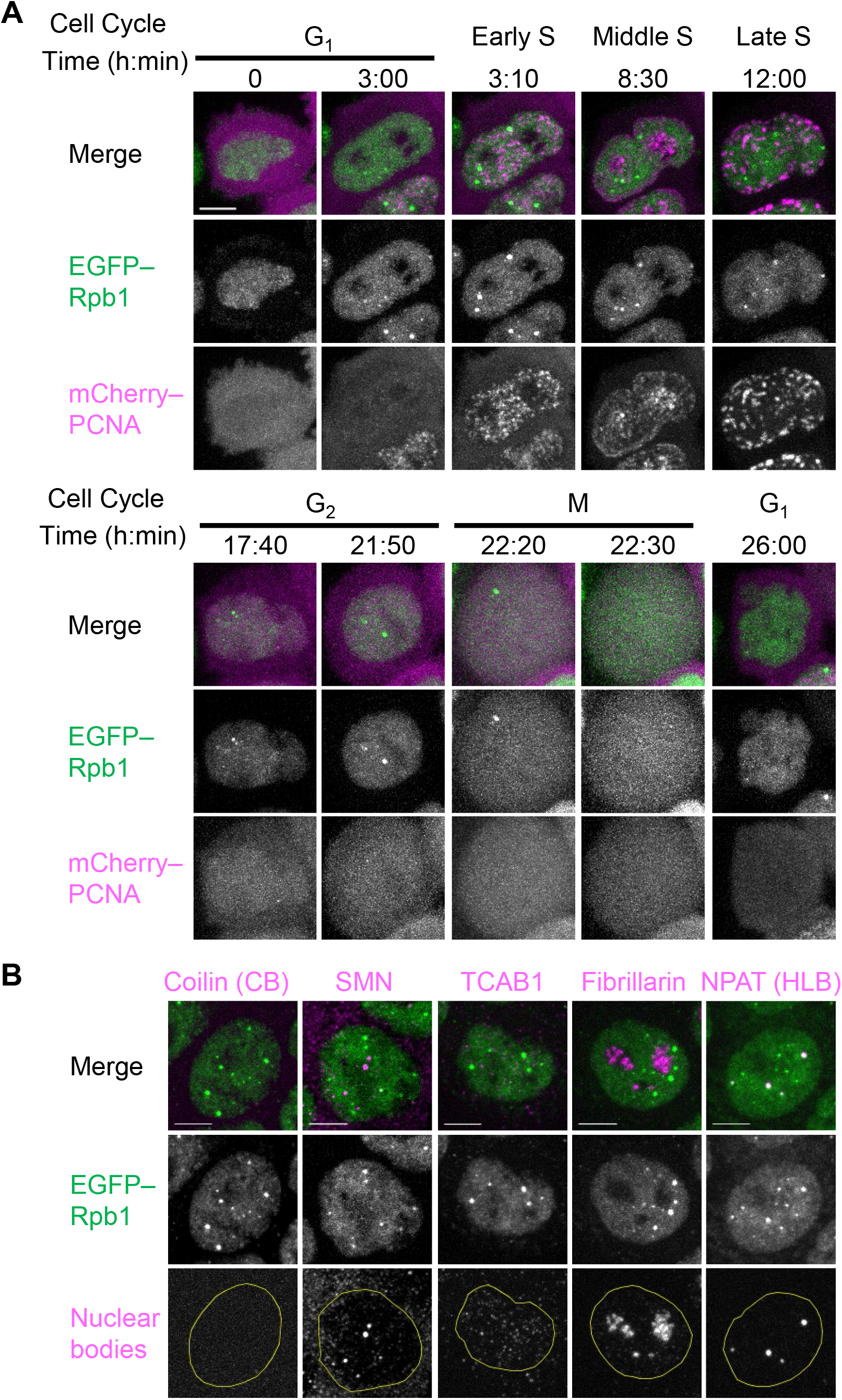
PCs in coilin-KO cells. **(A)** Coilin-KO cells that stably expressed mCherry–PCNA were established. Confocal images of EGFP–Rpb1 (green) and mCherry–PCNA (magenta) in the coilin-KO cells were acquired using time-lapse microscopy. MIP images of 15 Z-stacks at 0.75 μm intervals are shown. PCs nucleated in early S phase in coilin-KO cells, as observed in KI cells. **(B)** Coilin-KO cells were fixed and stained with antibodies against coilin, SMN, TCAB1, fibrillarin, and NPAT, and Cy5-labeled secondary antibodies (magenta). PCs were detected using EGFP–Rpb1 (green). Confocal images of the entire nucleus were acquired at 0.30 μm Z-intervals (29–62 stacks). MIP images are shown. Yellow lines indicate nuclear peripheries. Scale bars, 5 μm.

**Figure S6.**
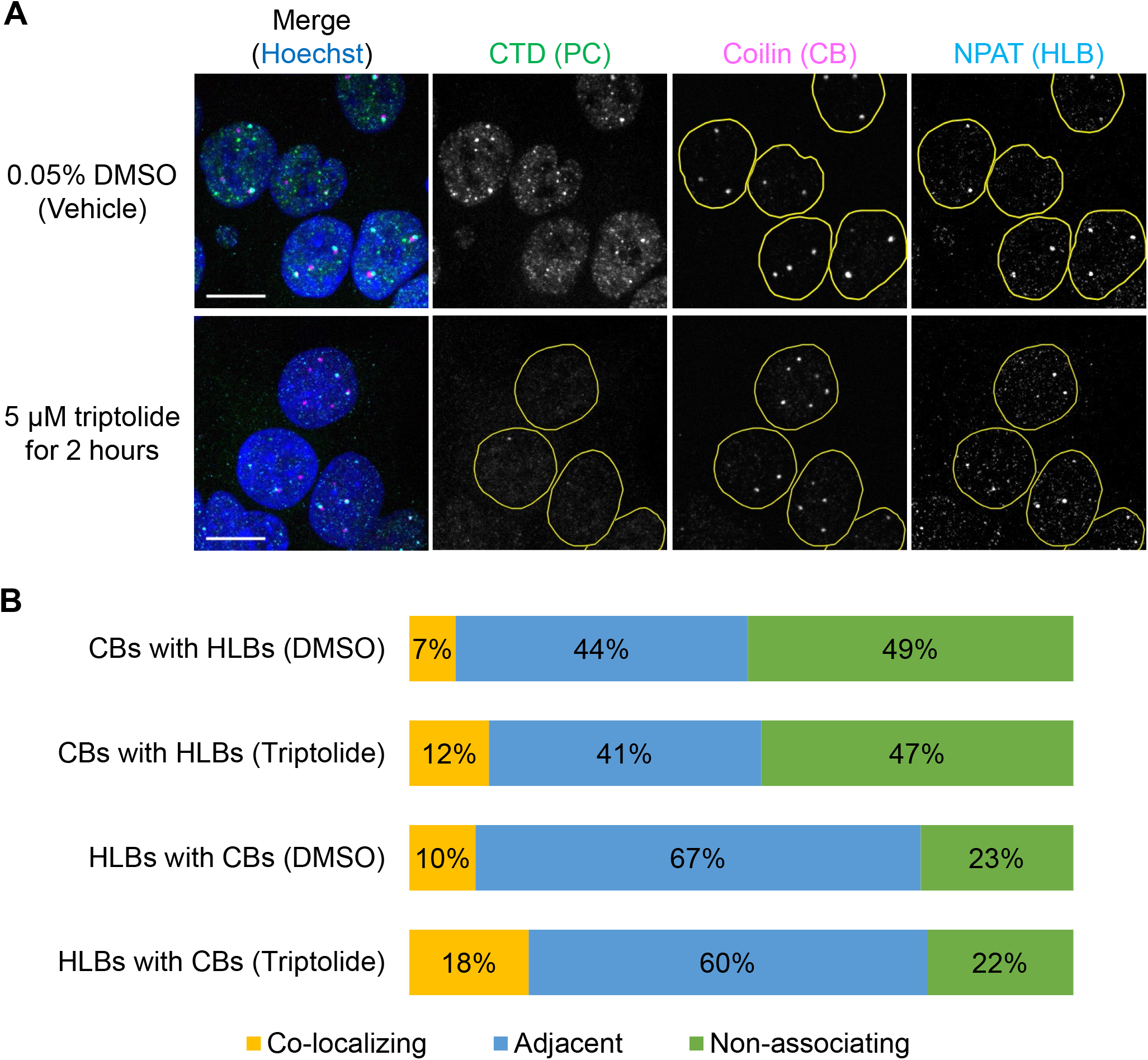
Effects of PC depletion by triptolide treatment on CB and HLB association in parental HCT116 cells. Parental HCT116 cells were treated with 5 μM triptolide or DMSO (vehicle) for 2 h, fixed, and stained with antibodies against Pol II CTD (PC; green), coilin (CB; magenta), and NPAT (HLB; cyan). Nuclear DNA was counterstained with Hoechst 33342 (blue). **(A)** MIP images of confocal sections. Confocal images of the entire nucleus were acquired at 0.30 μm Z-intervals (41–50 stacks). Yellow lines indicate nuclear peripheries. Scale bar, 10 μm. **(B)** Association rates between CBs and HLBs. “CBs with HLBs” indicates the percentage of CBs that are colocalized with, adjacent to, or non-associated with HLBs (n = 30 cells). “HLBs with CBs” indicates the percentage of HLBs that are colocalized with, adjacent to, or non-associated with CBs (n = 30 cells).

